# Multi-omics analysis of xylem sap uncovers dynamic modulation of poplar defenses by ammonium and nitrate

**DOI:** 10.1101/2021.05.28.446139

**Authors:** Karl Kasper, Ilka N Abreu, Kirstin Feussner, Krzysztof Zienkiewicz, Cornelia Herrfurth, Till Ischebeck, Dennis Janz, Andrzej Majcherczyk, Kerstin Schmitt, Oliver Valerius, Gerhard H. Braus, Ivo Feussner, Andrea Polle

## Abstract

Xylem sap is the major transport route for nutrients from roots to shoots. Here, we investigated how variations in nitrogen (N) nutrition affected the metabolome and proteome of xylem sap, growth of the xylem endophyte *Brennaria salicis* and report transcriptional re-wiring of leaf defenses in poplar (*Populus* x *canescens*). We supplied poplars with high, intermediate or low concentrations of ammonium or nitrate. We identified 288 unique proteins in xylem sap. About 85% of the xylem sap proteins were shared among ammonium- and nitrate-supplied plants. The number of proteins increased with increasing N supply but the major functional categories (catabolic processes, cell wall-related enzymes, defense) were unaffected. Ammonium nutrition caused higher abundances of amino acids and carbohydrates, while nitrate caused higher malate levels in xylem sap. Pipecolic acid and *N*-hydroxy-pipecolic acid increased whereas salicylic acid and jasmonoyl-isoleucine decreased with increasing N nutrition. Untargeted metabolome analyses revealed 2179 features in xylem sap, of which 863 were differentially affected by N treatments. We identified 122 metabolites, mainly from specialized metabolism of the groups of salicinoids, phenylpropanoids, phenolics, flavonoids, and benzoates. Their abundances increased with decreasing N. Endophyte growth was stimulated in xylem sap of high N- and suppressed in that of low N-fed plants. The drastic changes in xylem sap composition caused massive changes in the transcriptional landscape of leaves and recruited defense pathways against leaf feeding insects and biotrophic fungi, mainly under low nitrate. Our study uncovers unexpected complexity and variability of xylem composition with consequences for plant defenses.

**Significance statement:** This study reports the largest, currently available plant xylem sap proteome and metabolome databases and highlights novel discoveries of specialized metabolites and phytohormones in the xylem sap. This is the first multi-omics study linking differences in nitrogen supply with changes xylem sap composition, endophyte growth and transcriptional defenses in leaves.

## Introduction

Nitrogen (N) is a key plant nutrient, required for biosynthesis of amino acids, proteins, nucleic acids, and for many other essential metabolic functions. In soil, the dominant forms available for plant nutrition are nitrate and ammonium (Marschner, 2011). Their concentrations in the environment are highly variable, depending for example on soil type, microbial N turn-over, and atmospheric deposition (Nie *et al.*, 2017). In undisturbed forests, N is often a limiting factor for tree growth, requiring N cycling within the ecosystem (Schimel and Bennett, 2004); Rennenberg & Dannenmann, 2015). Degradation and mineralization of organically bound N result in the oxidized form NO_3_^−^, which is then reduced to ammonium (NH_4_^+^) by microbial activities; NH_4_^+^ is, thus, generally the main N source for forest trees, whereas in ruderal and riparian sites, NO_3_^−^ is the major form taken up by trees (Rennenberg *et al.*, 2009). Anthropogenic activities, especially intense crop and livestock production have increased N deposition across Europe (Morseletto, 2019; Stevens, 2019). Large-scale N deposition has also increased in many forest regions (Rennenberg & Dannenmann, 2015). Excess N deposition results in an enhanced release of the greenhouse gas N_2_O to the atmosphere (Merbach *et al.*, 1996) and enhances NO_3_^−^ input and soil acidification with adverse effects on tree health and forest biodiversity (Jandl *et al.*, 2012). It is, therefore, important to better understanding how tree species cope with varying N availabilities in the environment.

Nitrogen is taken up by plant roots, metabolized and transported via the xylem sap to the leaves for further metabolization (Marschner, 2011). The consequences of variations in nitrate or ammonium nutrition have been studied in leaves and roots showing, for example, stronger responses to ammonium than to nitrate depletion in Arabidopsis leaves (Menz *et al.*, 2016). N transport in xylem sap has been studied in tree species focusing mainly on amino acids, nitrate, and ammonium (e.g., Gessler et al., 1998, 2004; Kato, 1981; Siebrecht & Tischner, 1999; Tromp & Ovaa, 1976; Weber et al., 1998). Poplar (*Populus* x *canescens*) xylem sap contains approximately 3 mM nitrate and up to 15 mM glutamine, when fed with 1 mM NO_3_^−^ in hydroponics (Siebrecht and Tischner, 1999). After switching from nitrate to ammonium nutrition, nitrate concentrations in the xylem sap of poplar drop quickly and NH_4_^+^ levels increase (Siebrecht and Tischner, 1999). In addition to mineral nutrients and amino acids, xylem sap also contains a huge array of proteins with roles in cell wall formation and defense (Dafoe and Constabel, 2009; Pechanova *et al.*, 2010). However, comprehensive analyses, if and how changes in ammonium or nitrate availabilities influence the metabolome and proteome of the xylem sap are lacking.

Ecological studies reported that N nutrition affects the balance between growth and defense, thereby, interfering with biotic stress tolerance (Harding *et al.*, 2009; Manninen *et al.*, 1998; Nerg *et al.*, 2008; Rubert-Nason *et al.*, 2015). For example, high N availability results in decreased tannin concentrations in poplar leaves (Madritch and Lindroth, 2015) and enhanced development of the leaf feeding larvae of *Choristoneura conflictana* (Bryant *et al.*, 1987). In various agricultural crops, infections with fungal pathogens decreased under low and increased under high N supply (Mur *et al.*, 2017), suggesting that information conveyed by the xylem sap from roots to leaves has consequences for foliar defense systems.

The main aims of the present work were a comprehensive characterization of the metabolome and proteome of *P.* x *canescens* xylem sap in response to increasing N supplied either as nitrate or as ammonium. A further important goal was to investigate the responses of the leaf defense transcriptome to changes in N supply. For this purpose, *P.* x *canescens* were supplied with low (0.4 mM), medium (2 mM), or high concentrations (8 mM) of NH_4_^+^ or NO_3_^−^. The composition of the xylem sap in response to these changes in N supply was extensively characterized, resulting in the largest libraries for poplar xylem sap metabolome and proteome data available to date. We report for the first time pipecolic acid and *N*-hydroxy-pipecolic acid (NHP) in xylem sap and their differences under ammonium and nitrate nutrition. Changes in nitrate and ammonium concentrations also had divergent effects on the xylem sap composition of amino acids, carbohydrates, organic acids, secondary compounds, and phytohormones. Protein diversity in the xylem sap increased with increasing N supply. We tested the impact of these changes for poplar susceptibility to biotic stress in the vascular system employing bioassays with the gram-negative bacterium *Brennaria salicis* (formerly *Erwinia salicis*). *B*. *salicis* lives as an endophyte in the xylem of *Populus spp*. (Maes *et al.*, 2009) and is the causal agent of the watermark disease in *Salix* spp. (Day, 1924; Huvenne *et al.*, 2009). By bioinformatic comparison of poplar leaf transcriptomes against poplar rust (*Melamspora* spp., Luo et al., 2019; Miranda et al., 2007; Rinaldi et al., 2007) and herbivory (*Chrysomela populi,* Kaling et al., 2018), we found low-nitrate driven activation of salicylic acid (SA)- and jasmonate (JA)-related defenses.

## Results

### Ammonium and nitrate differentially shape primary metabolite composition in xylem sap

Feeding poplars with nitrate or ammonium strongly affected the concentrations of these nutrients in poplar xylem sap (Table 1). The concentrations of NO_3_^−^ increased with increasing nitrate and of NH_4_^+^ with increasing ammonium supply (Table 1). The highest protein concentrations occurred in xylem sap of plants fed with high ammonium (Table 1). In agreement with other poplar studies (Euring *et al.*, 2014; Gan *et al.*, 2015; Reichardt *et al.*, 1991), increasing N supply resulted in enhanced photosynthesis, growth, biomass, and increased tissue concentrations of N (Supporting Table S1, Supporting Figure S1).

**Table 1:** Xylem sap NH_4_-N, NO_3_-N, and protein concentrations in *Populus* x *canescens* treated with different forms and concentrations of nitrogen

| Treatment | Xylem sap $\text{NH}_4\text{-N}$<br>(mM) | Xylem sap $\text{NO}_3\text{-N}$<br>(mM) | Xylem sap protein<br>($\mu\text{g mL}^{-1}$) |
| --- | --- | --- | --- |
| $\text{NH}_4^+$ L | 8.5 ± 4.8 a | 2.7 ± 1.0 a | 3.7 ± 1.4 a |
| $\text{NH}_4^+$ M | 33.5 ± 10.7 a | 1.6 ± 0.2 a | 5.7 ± 1.8 a |
| $\text{NH}_4^+$ H | 210.8 ± 33.2 b† | 1.2 ± 0.2 a | 23.7 ± 2.5 c |
| $\text{NO}_3^-$ L | 7.1 ± 2.9 a | 3.3 ± 1.0 a | 5 ± 0.5 a |
| $\text{NO}_3^-$ M | 26.3 ± 7.0 a | 81.2 ± 92.0 a | 5 ± 1.0 a |
| $\text{NO}_3^-$ H | 29 ± 18.4 a | 386.4 ± 268.5 b† | 9.8 ± 2.8 b |

L = 0.4 mM, M = 2.0 mM, H = 8.0 mM nitrogen supplied in nutrient solution. Data show means ( $n=5 \pm \text{SD}$ ), unless indicated by a †. Those data are means of $n=10 (\pm \text{SD})$ . Different letters in a column indicate significant differences at $p \leq 0.05$ according to Tukey's-Kramer post-hoc test.

Intermediate and high ammonium nutrition further resulted in high amino acid concentrations in the xylem sap, whereas the increases in amino acids were less pronounced under high nitrate nutrition (Figure 1a, Supporting Table S2). The main amino acids detected under the current measuring conditions were derived from glutamate. Since glutamine and glutamate spontaneously cyclized to pyroglutamate during GC-MS sample preparation (Supporting Figure S2), the contributions of those individual compounds to xylem sap amino acids cannot be distinguished here. Taken together, glutamine and glutamate accumulated more than twelve-fold under high compared to low ammonium supply but only up to six-fold under high compared to low nitrate supply. Other studies found that glutamine is the major amino acid in poplar xylem sap (Dickson et al., 1985; Siebrecht & Tischner, 1999).

**Figure 1:**
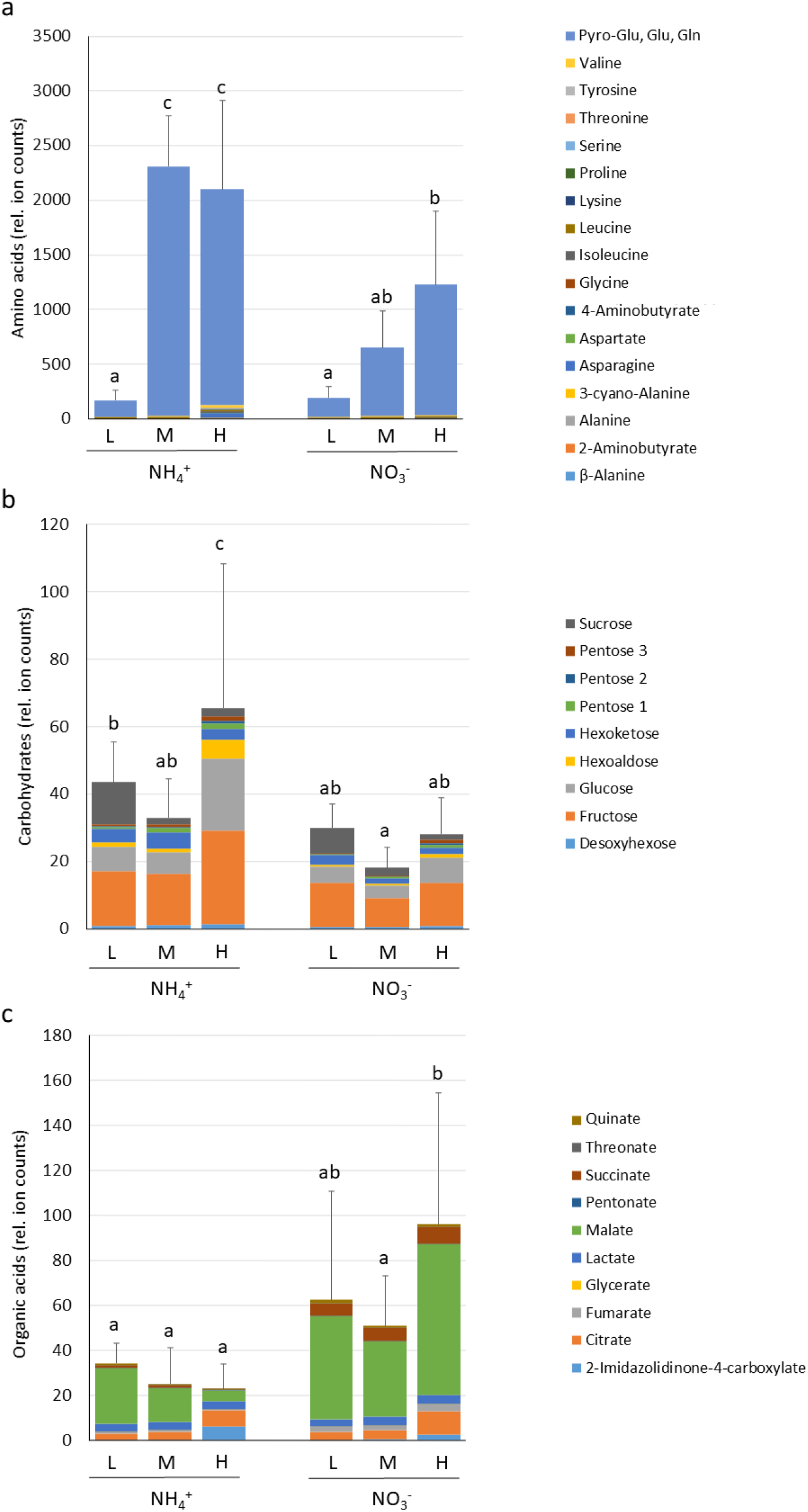
Relative abundances of a) amino acids, b) soluble carbohydrates and c) organic acids in *Populus* x *canescens* xylem sap. Xylem saps were collected three weeks after feeding poplars with L (0.4 mM), M (2.0 mM) or H (8.0 mM) ammonium or nitrate in the nutrient solution. Pyr-Glu, Glu, Gln indicates the sum of pyroglutamate, glutamate and glutamine. Supportive Figure S2 provides further information. Stacked bars show means of the relative abundances for the indicated compounds. Error bars refer to the variation of the sum of compounds. Different letters above the bars indicate significant differences at p ≤ 0.05 according to Tukey’s post-hoc test, n=8 per treatment.

Furthermore, ammonium and nitrate nutrition caused alterations in carbon-bearing compounds in the xylem sap: soluble carbohydrate levels were approximately twofold higher in high ammonium- than in high nitrate-fed plants (Figure 1b). Soluble organic acids were generally higher under nitrate than under ammonium nutrition, resulting in approximately five-fold increases in xylem sap of high nitrate-compared to that of high ammonium-fed plants (Figure 1c). Fructose and glucose were the most prominent carbohydrates and malate the most abundant organic acid in xylem sap (Figure 1 b,c).

### Ammonium- and nitrate-feeding impact phytohormones in the xylem sap

We detected SA, SA-glucoside (SAG), jasmonoyl-isoleucine (JA-Ile), pipecolic acid, NHP, and abscisic acid (ABA) in poplar xylem sap (Figure 2). SA, SAG, JA-Ile, pipecolic acid and NHP are involved in regulating plant defense responses (Antico *et al.*, 2012; Bernsdorff *et al.*, 2016; Y.,-C., Chen *et al.*, 2018; Schuman *et al.*, 2018; Vlot *et al.*, 2009). Xylem sap of low ammonium-fed poplars contained the highest concentrations of SA, SAG, and JA-Ile (Figure 2a,b,c). The concentrations of these phytohormones decreased with increasing N concentrations and were generally higher in ammonium-than in nitrate-fed plants. On the contrary, pipecolic acid and NHP increased with increasing NH_4_^+^ supply (Figure 2d,e). Xylem sap of high ammonium-fed poplars contained the highest pipecolic acid and NHP levels, whereas NHP was not significantly affected by low and medium, and not detected under high nitrate supply (Figure 2d,e). ABA, which is important to coordinate the plant’s stature and acclimation to drought (Fujita *et al.*, 2005; Yu *et al.*, 2019), was unaffected by differences in N supply in poplar xylem sap (Figure 2f).

**Figure 2:**
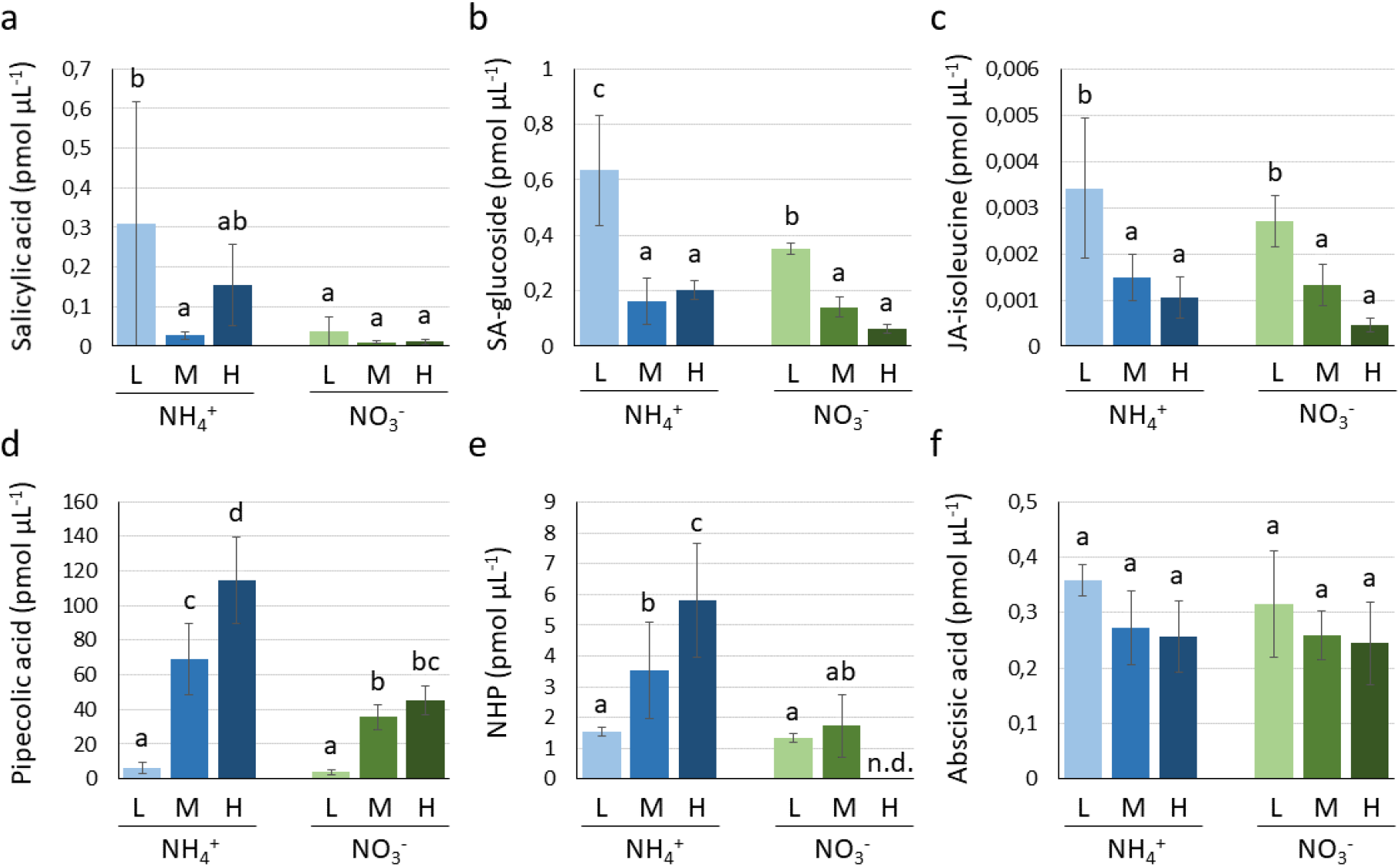
Phytohormone concentrations in *Populus* x *canescens* xylem saps. a) Salicylic acid (SA), b) SA-glucoside (SAG), c) Jasmonoyl-isoleucine (JA-Ile), d) Pipecolic acid, e) *N*-hydroxypipecolic acid (NHP), f) Abscisic acid (ABA). Additional phytohormones identified in xylem sap are listed in supplemental Table S3. Xylem saps were collected three weeks after feeding poplar with L (0.4 mM), M (2.0 mM) or H (8.0 mM) ammonium or nitrate in the nutrient solution. n.d. = not detected. Data were log_2_ transformed to achieve homogeneity of variance for statistical analyses. Bars show means (± SD) per treatment. Different letters above the bars indicate significant differences at p ≤ 0.05 according to Tukey’s-Kramer post-hoc test, n=5 to 6 per treatment.

### Defense metabolites increase in poplar xylem sap under low nitrogen supply

Next, we aimed to obtain a global overview on changes of xylem sap metabolites in response to N nutrition of poplar using a non-targeted metabolome approach. Alterations in the abundance of specialized metabolites, which had not been covered by the analyses shown so far, were of special interest in this experiment. In total, 2179 metabolite features were obtained by the non-targeted approach, a number that was reduced to 863 by filtering the data by a false discovery rate FDR < 10^−6^ (Supportive data file S1, Supporting Figure S3). A sample-based principal component analysis (PCA) for these selected features discriminated the samples along PC1 according to the N concentrations and along PC2 according to the N form (ammonium or nitrate) supplied to the plants (Figure 3a). Feature-based clustering by one-dimensional self-organizing maps was used for a general overview on the intensity patterns of the selected features (Supporting Figure S4). The heat map representation of the data in the self-organizing maps showed that each of the six nutrient treatments was characterized by a specific response pattern of metabolite features in the xylem sap, depending on the N form as well as on the N concentration supplied to the plants (Supporting Figure S4). The heat map indicated that a large number of features increased with decreasing N supply, irrespective of the N form, whereas the number of features strongly increasing with increasing N supply was higher in xylem sap of ammonium (about 180 features) than in that of nitrate-fed poplars (12 features, Supporting Figure S4).

**Figure 3:**
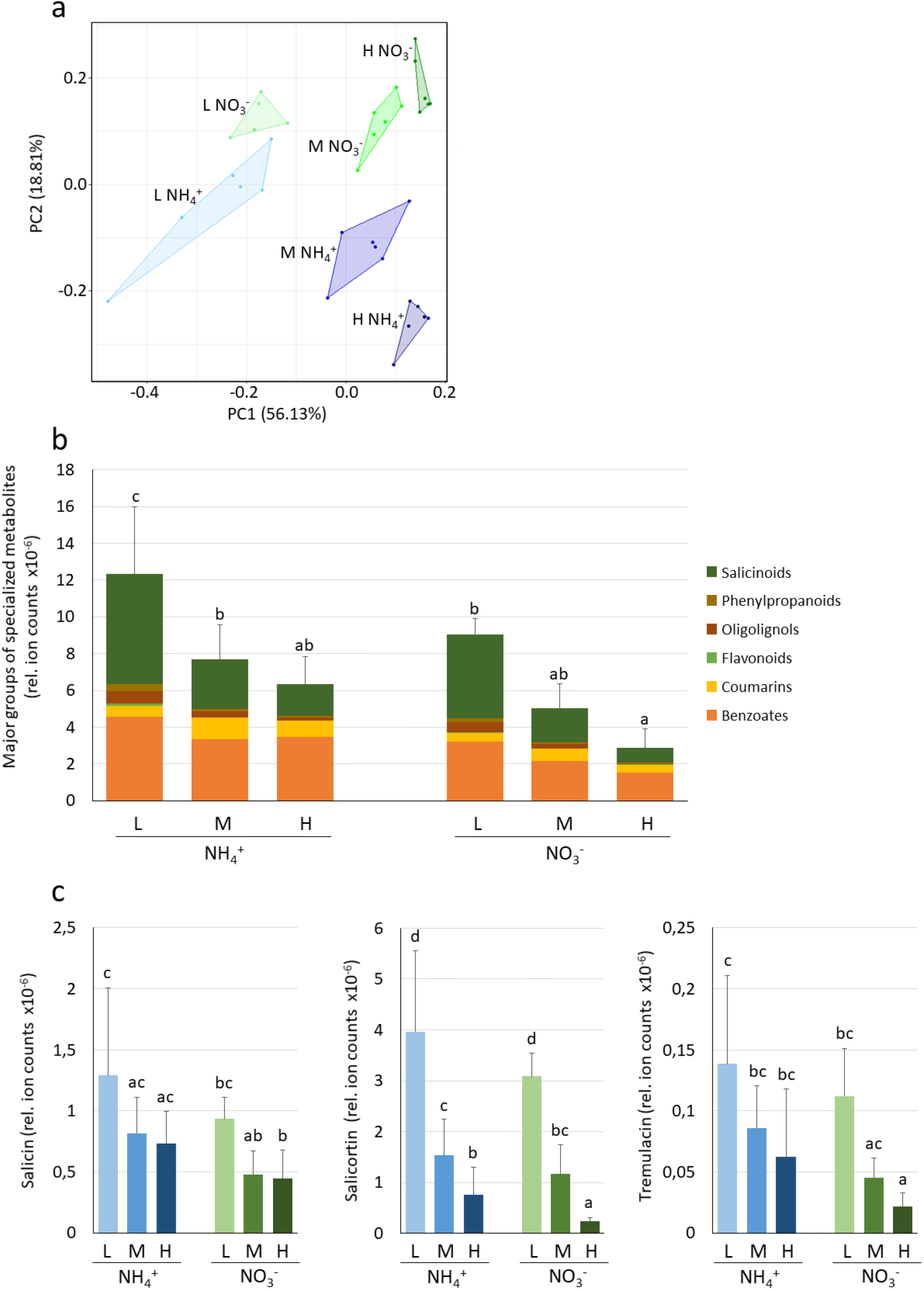
Specialized metabolite profiles in *Populus* x *canescens* xylem saps. a) Principal component analysis of 863 metabolite features of a non-targeted metabolome approach (metabolite fingerprinting) with FDR < 10^−6^, b) Relative signal intensities of the major specialized metabolite classes, c) Relative signal intensities of three major salicinoid compounds. Xylem saps were collected three weeks after feeding poplar with L (0.4 mM), M (2.0 mM) or H (8.0 mM) ammonium or nitrate in the nutrient solution. Data were log_2_ transformed to achieve homogeneity of variance for statistical analyses and show means (± SD) per treatment of five to six biological replicates. Compounds present two independent experiments (Supportive Figure S3) were included. Different letters above the bars indicate significant differences at p ≤ 0.05 according to Tukeys-Kramer post-hoc test.

To determine the chemical identity of the compounds, represented by the features in the data set, the accurate mass information of the features was used for data base search and LC-HRMS/MS fragmentation analyses were conducted. By this approach, the identities of 75 specialized metabolites were determined in the xylem sap of poplar (Supportive data file S2). Compounds involved in poplar defense (Boeckler *et al.*, 2011; Chedgy *et al.*, 2015), like salicinoids, phenylpropanoids, flavonoids, and benzoates (Figure 3b) were identified. Among the identified classes of secondary metabolites, salicinoids and benzoates accounted for the highest accumulated signal intensities (Figure 3b). Overall, the relative abundances of the specialized metabolites decreased with increasing N supply, with the exception of coumarins, which were differently affected by N treatments (Figure 3b) and exhibited lowest values under low NH_4_^+^ and all NO_3_^−^ treatments (p < 0.001). Among the salicinoids, salicortin showed high signal abundance under low N supply, which declined to one fifth after high N supply (Figure 3c). Also, salicin and tremulacin showed the highest abundances in xylem sap of poplars fed with low N (Figure 3c).

With the number of 75 identified compounds, a substantial fraction of metabolites represented by the data set was determined, considering the large number of adducts known to be formed by ESI-MS. Our analyses also confirmed metabolites already detected by the targeted approaches (Figure 1 and 2), like organic acids, amino acids but also ABA, pipecolic acid and NHP (Supportive data file S2). Overall, the combined approaches of targeted primary metabolite analysis, phytohormone quantification and non-targeted LC-MS metabolite fingerprinting with subsequent MS/MS based metabolite annotation led to the identification of 122 metabolites and, thus, is the largest xylem sap metabolite database currently available (Supporting Table S3).

### Increasing nitrogen concentrations affect the xylem sap proteome diversity

Proteome analyses showed the highest protein diversity in the xylem sap of high ammonium- and high nitrate-fed plants (Supportive data file S3), indicating that not only the protein concentrations (Table 1) but also the number of proteins increased (Figure 4a). A high number of the xylem sap proteins had a signal peptide, targeting the compound to the extracellular compartment (62.5%) (Supportive data file S3). Since we did not detect any significant contamination of the xylem sap with the intracellular marker enzyme glucose-6-phophate dehydrogenase (cf. Experimental procedures) and since proteins can also be targeted by other mechanisms to the apoplast (Feussner & Polle, 2015), proteins without known apoplastic target sequence were also considered as likely xylem sap constituents. Rare proteins might have escaped the label-free quantification approach (Cox *et al.*, 2014).

**Figure 4:**
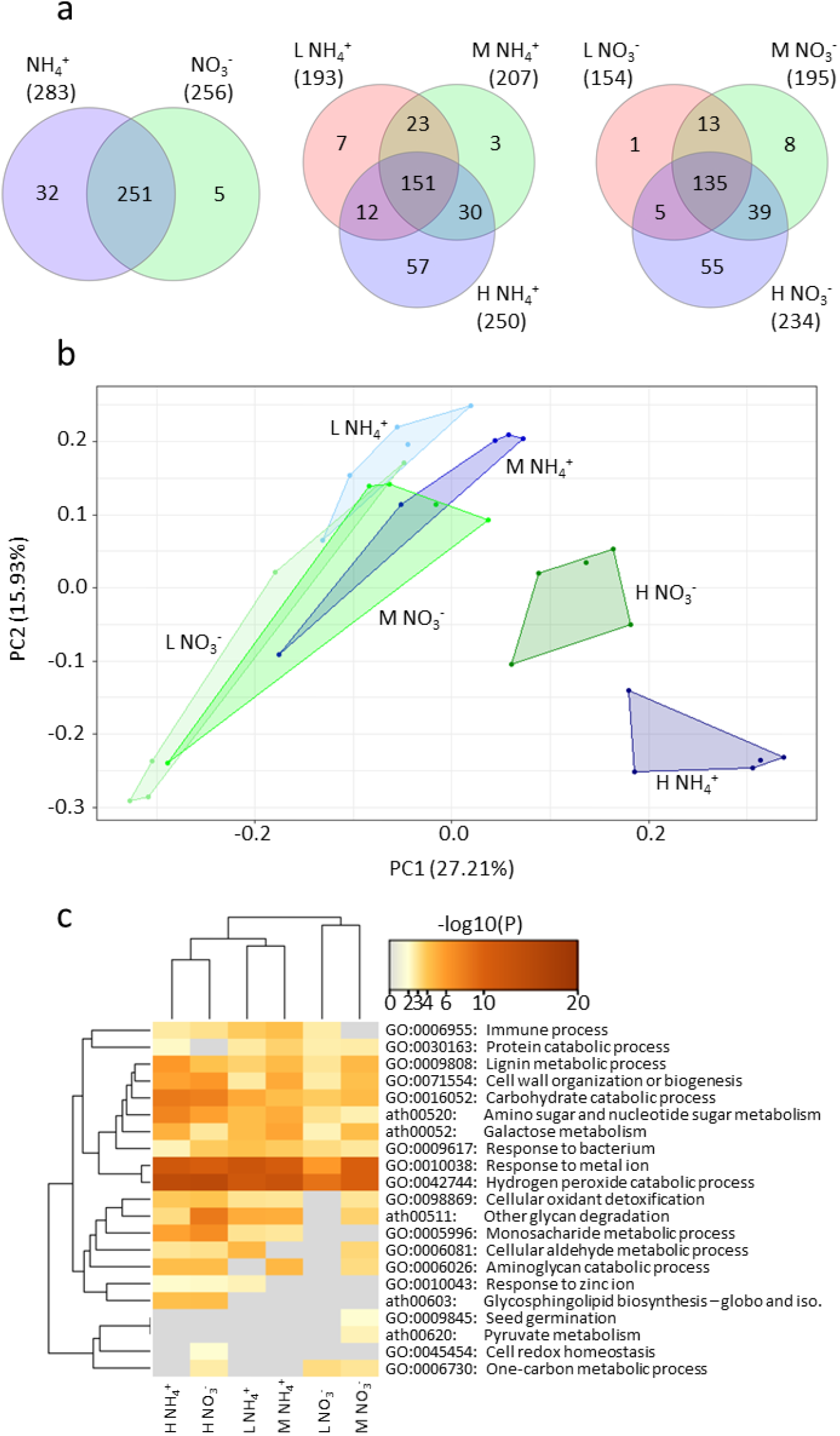
Proteome composition in *Populus* x *canescens* xylem saps. a) Venn diagrams showing numbers of unique and overlapping proteins in xylem sap of plants supplied with supplied with low (L), intermediate (M) or high (H) NH_4_^+^ or NO_3_^−^, b) Principal component analysis of the proteins in the xylem sap from different N treatments, c) Hierarchical cluster analysis of significantly enriched gene ontology terms and KEGG pathways of poplar xylem sap proteins. Analyses were conducted with the best AGI matches of the poplar proteins and standard settings in Metascape. Xylem saps were collected three weeks after feeding poplar with L (0.4 mM), M (2.0 mM) or H (8.0 mM) ammonium or nitrate in the nutrient solution. Proteins present in 10 or more percent of samples were used for analyses. n=5 per treatment.

Filtering the proteins that could be quantified in ≥ 10% of the samples resulted 289 unique proteins, of which 288 were assigned to a poplar gene identifier (Potri ID) and could be mapped to 222 unique Arabidopsis AGIs (Supportive data file S3). The overlap between the ammonium and nitrate treatments was high with 251 shared poplar proteins (Figure 4a).

Under ammonium 32 and under nitrate nutrition five unique proteins were detected (Figure 4a). PCA showed strong overlap of the xylem sap protein profiles obtained under low and intermediate ammonium and nitrate nutrition, whereas xylem sap proteins of high ammonium- and high nitrate-fed plants were clearly separated (Figure 4b).

GO term and KEGG pathway analysis of the xylem sap proteins on the Metascape platform revealed 21 significantly enriched biological functions (Figure 4c). Major functional classes were defense (“Immune response”, “Hydrogen peroxide degradation”, and “Cellular oxidant detoxification”), cell wall-related processes (“Lignin metabolic process”, “Cell wall organization” and “Carbohydrate metabolic processes”) and catabolic processes (“Monosaccharide metabolic process”, “Pyruvate and aldehyde metabolism”, “Aminoglucan metabolism”, “Amino sugar and nucleotide sugar metabolism”) (Figure 4c). The GO term “Immune response” comprised natriuretic protein, lysine-motif domain GPI-anchored protein 1 (LYM1), peroxidases, chitinases, and bifunctional inhibitor/lipid-transfer protein/seed storage 2S albumin superfamily protein. The GO terms “Cell wall metabolism” and “Lignification” included xylanases, glucanases, peroxidases, and laccases and suggested functions such as pectin and hemicellulose metabolism and catalytic activities driving the last step of lignification by H_2_O_2_-dependent peroxidases and O_2_-dependent laccases (Supportive data file S3). The fewest significant changes in GO terms occurred when the N supply was increased from low to intermediate ammonium or nitrate levels (Figure 4c).

Using DESeq2 (Love *et al.*, 2014) on protein abundance data (Supportive data file S3), we identified 59 xylem sap proteins that showed significant differences (P_adj_ < 0.05) in response to changes in N nutrition. The significant proteins in xylem sap from low and intermediate ammonium fed plants clustered together and were distinct from low and intermediate nitrate-fed plants (Figure 5). Proteins in xylem sap from high ammonium- and high nitrate-fed plants formed distinct clusters with the highest number of differentially expressed protein. Under both high ammonium and high nitrate, similar sets of stress-related protein showed higher abundances than under lower N supply including putative chitinases, peroxidases, laccase, proteases, and peptidases (Figure 5). Overall, our analyses shows that proteins patterns in xylem sap responded strongly to changes in the N concentrations, especially in the category of phenylpropanoid metabolism. By identification of 289 proteins, we present the largest current xylem sap proteome in a tree species (Supportive data file S3).

**Figure 5:**
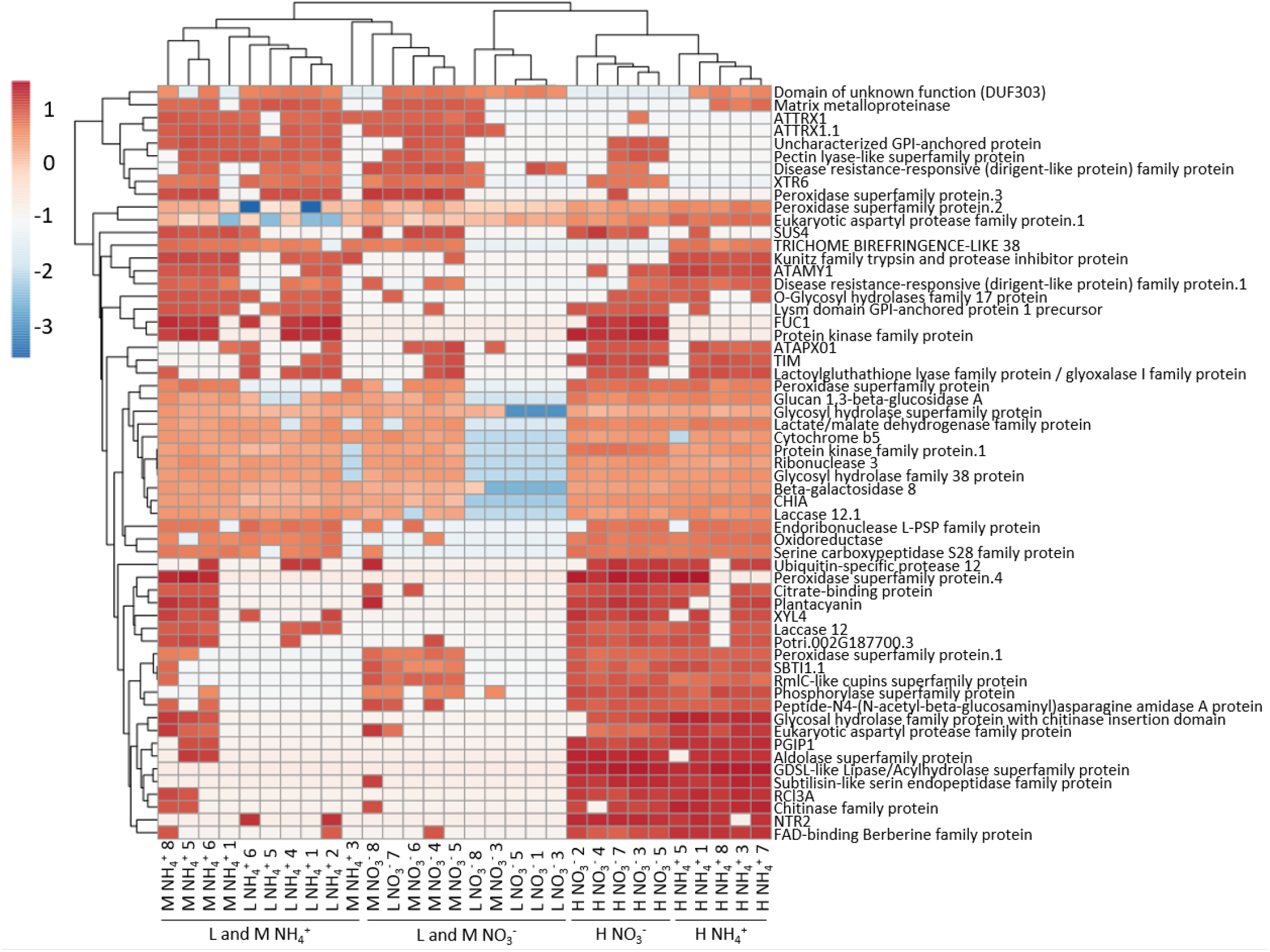
Hierarchical cluster analysis of 59 xylem sap proteins with significantly different abundances between nitrogen treatments. The analysis was generated with ClustVis and standard settings. *Populus* x *canescens* xylem saps were collected three weeks after feeding poplars with L (0.4 mM), M (2.0 mM) or H (8.0 mM) ammonium or nitrate in the nutrient solution. n=5 per treatment.

### High plant nitrogen availability stimulates growth of a poplar xylem endophyte

To investigate whether the compositional metabolic changes and shifts in the defense proteome affected xylem-colonizing bacteria, we studied growth of *B. salicis* using xylem sap as the growth medium. Bacterial growth was strongest in xylem sap of poplars fed with high ammonium (Figure 6). Xylem sap from plants with intermediate ammonium supply and high nitrate supply afforded similar growth of *B*. *salicis* (Figure 6). The lowest growth of *B. salicis* was found in xylem sap of poplars with low nitrate supply (Figure 6). Obviously, *B. salicis* benefited from an increased nutritional quality of xylem sap from poplars fed with high ammonium.

**Figure 6:**
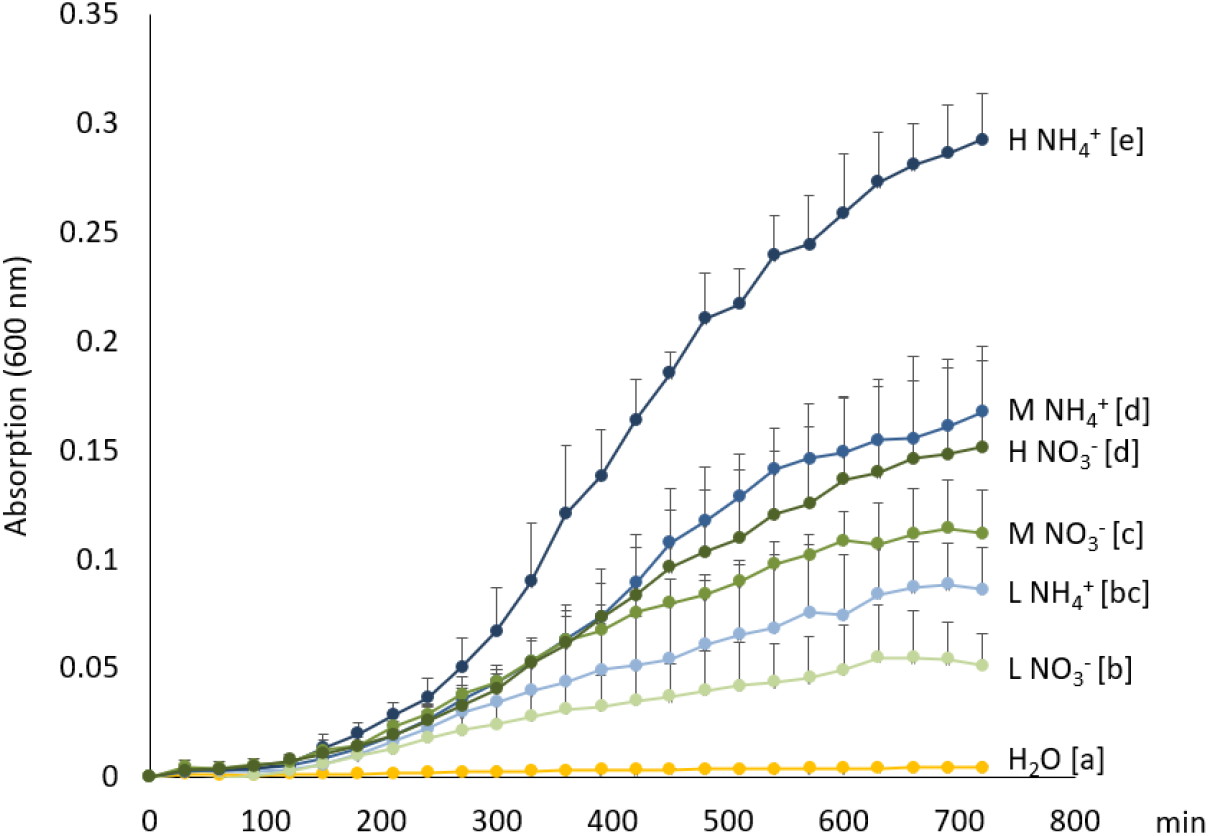
*Brennaria salicis* growth in xylem sap of *Populus* x *canescens*. *Populus* x *canescens* xylem saps were collected three weeks after feeding poplars with L (0.4 mM), M (2.0 mM) or H (8.0 mM) ammonium or nitrate in the nutrient solution. Aliquots of xylem sap of eight biological replicates per treatment were inoculated with *B. salicis* or water (control) and the increase in the optical density was determined for 12h. Different letters in () indicate significant differences at p ≤ 0.05 according to Tukey’s post-hoc test. Data are means (n=8 ±SD).

### High and low N supply elicit stress and immune responses in the leaf transcriptome

Changes in the N concentrations in the nutrient solution had drastic effects on the leaf transcriptomes resulting in 9157 differentially expressed genes (DEGs with more than 2-fold changes and p_adj_ < 0.05) in response to changes in N concentrations and 431 DEGs in response to ammonium versus nitrate nutrition (Supportive data file S4). Global nitrogen effects have already been analyzed in several studies (e.g. Robertson & Vitousek, 2009; Scheible et al., 2004) and therefore, further details are shown in the supplements (Supporting Figure S5). Here, we focused on the question whether our nitrogen treatments elicited distinct immune responses known to occur in response to leaf-feeding insects and methyl jasmonate (MeJA) or in response to biotrophic fungi and SA treatments (Pieterse *et al.*, 2014).

To test whether changes in N nutrition resulted in MeJA/herbivory or SA/pathogen-related responses, we applied an *in silico* strategy: we extracted DEGs (> 2-fold change, p_adj_ < 0.05) in response to MeJA, to SA (Luo *et al.*, 2019), to rust fungi (*Melampsora medusae,* Miranda *et al.,* 2007; *Melampsora larici-populina,* Rinaldi *et al.,* 2007; Luo *et al.,* 2019), and to herbivory by poplar leaf beetle (*Chrysomela populi,* Kaling *et al.,* 2018) from previous publications.

We combined the MeJA/Chrysomela (MeJA+Chry) data sets keeping only genes that showed the same direction of change in response to MeJA and in response to Chrysomela (Supportive data file S5). Then, the transcript abundances of the genes in our N data set that overlapped with the MeJA+Chry matrix were extracted (Supportive data file S5) and clustered in a heatmap (Fig. 7a). DEGs regulated by MeJA and beetle feeding exhibited five main clusters (113 genes) that overlapped with N-responsive genes (Figure 7a). Cluster 1 contained genes with enhanced transcript abundances under low nitrate compared with other N-treatments (Figure 7a). Under MeJA exposure as well as under Chrysomela feeding, the majority of the genes in cluster 1 showed enhanced transcript abundances (see left hand color code in Figure 7a). The genes in cluster 1 were functionally annotated as “JA-mediated signaling pathway”, “Response to lipid” and “Wounding” (Figure 7b,c). The genes in cluster 4 and 5 showed highest transcript abundances under high N supply, especially in response to ammonium (cluster 4), were in most cases positively related to MeJA+Chry induced genes (Figure 7a), and annotated as “Response to wounding”, “Response to water deprivation”, “Defense response”, and “Drug catabolic process” (Figure 7b,c). Cluster 2 and 3 contained genes with decreased transcript abundances under high N nutrition. The majority of these genes also showed negative responses to MeJA+Chry (Figure 7a). Cluster 2 and 3 were functionally annotated as “Cell wall organization”, “Anatomical structure”, and “Cellular lipid metabolic process” (Figure 7b,c).

**Figure 7:**
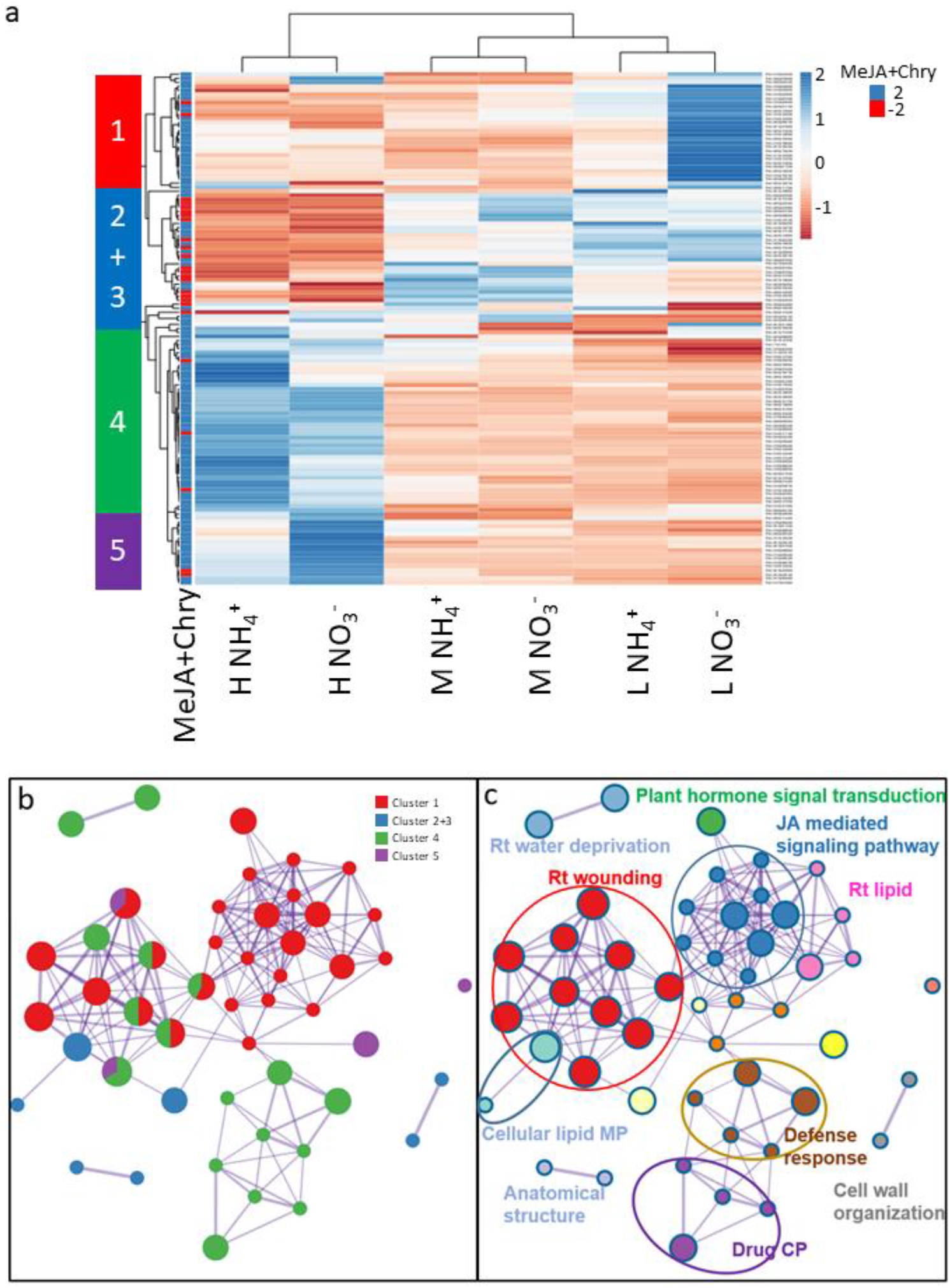
Cluster and network analyses of a set of shared differentially expressed genes (DEGs) in leaves of *Populus* x *canescens* exposed to different nitrogen and DEGs responding to application of methyl jasmonate (MeJA) and herbivory. DEGs for MeJA and herbivory by poplar leaf beetle (*Chrysomela populi* = Chry) were extracted from published studies (Kaling et al., 2018; Luo et al., 2019). a) hierarchical cluster analysis visualizing the pattern of the N-responsive DEGs. MeJA+Chry indicate positive (blue) or negative (red) response of DEGs to MeJA treatment or herbivory. The stacked bar on the left-hand side shows a color code for the five main clusters, b) enrichment network of DEGs within clusters of same color, c) biological processes assigned to DEGs shown in b. Leaves for transcriptome analyses were collected three weeks after feeding poplar with Low (0.4 mM), Medium (2.0 mM) or High (8.0 mM) ammonium or nitrate in the nutrient solution. Bioinformatic analysis were conducted with the best AGI match to the poplar IDs and the standard settings in Metascape. CP = catabolic process, JA = jasmonic acid, MP = metabolic process, Rt = response to. n=4 to 5 per treatment.

We employed the same strategy to identify leaf rust (*Melampsora* spp.) and SA co-regulated genes (SA+Mel), their overlap with N-responsive genes, and clustered the resulting set of 219 genes according to N-treatments (Fig. 8a, Supportive data file S5). Cluster 1 contained genes with enhanced transcript abundances under high nitrate that were functionally annotated as “Ethylene activated signaling” (Fig. 8b,c). Cluster 2 contained genes with high transcript abundances for high ammonium and nitrate treatments encompassing the categories “Organic anion transport”, “Response to oxidative stress” and “Cell wall biogenesis” (Figure 8b,c). Genes in cluster 4, which showed high transcript abundances under low nitrate treatments, were functionally annotated as “Hormone biosynthetic process” and “Secondary metabolism” (Figure 8a-c). Cluster 5 showed high transcript abundances for low nitrate-fed plants (Figure 8a). Cluster 5 was the largest one, with “Defense response to fungus”, “Response to bacterium”,” Immune system process” and “Regulation of hormone metabolic process” (Figure 8b,c). In most cases, the genes in cluster 4 and 5 were positively related to SA+Mel induced genes (Figure 8a). By our comparison of N-, biotic stress- and phytohormone-responsive DEGs, we provide a novel resource that can be used in the future to deepen our understanding of the relationship between N supply and tree stress susceptibility or tolerance.

**Figure 8:**
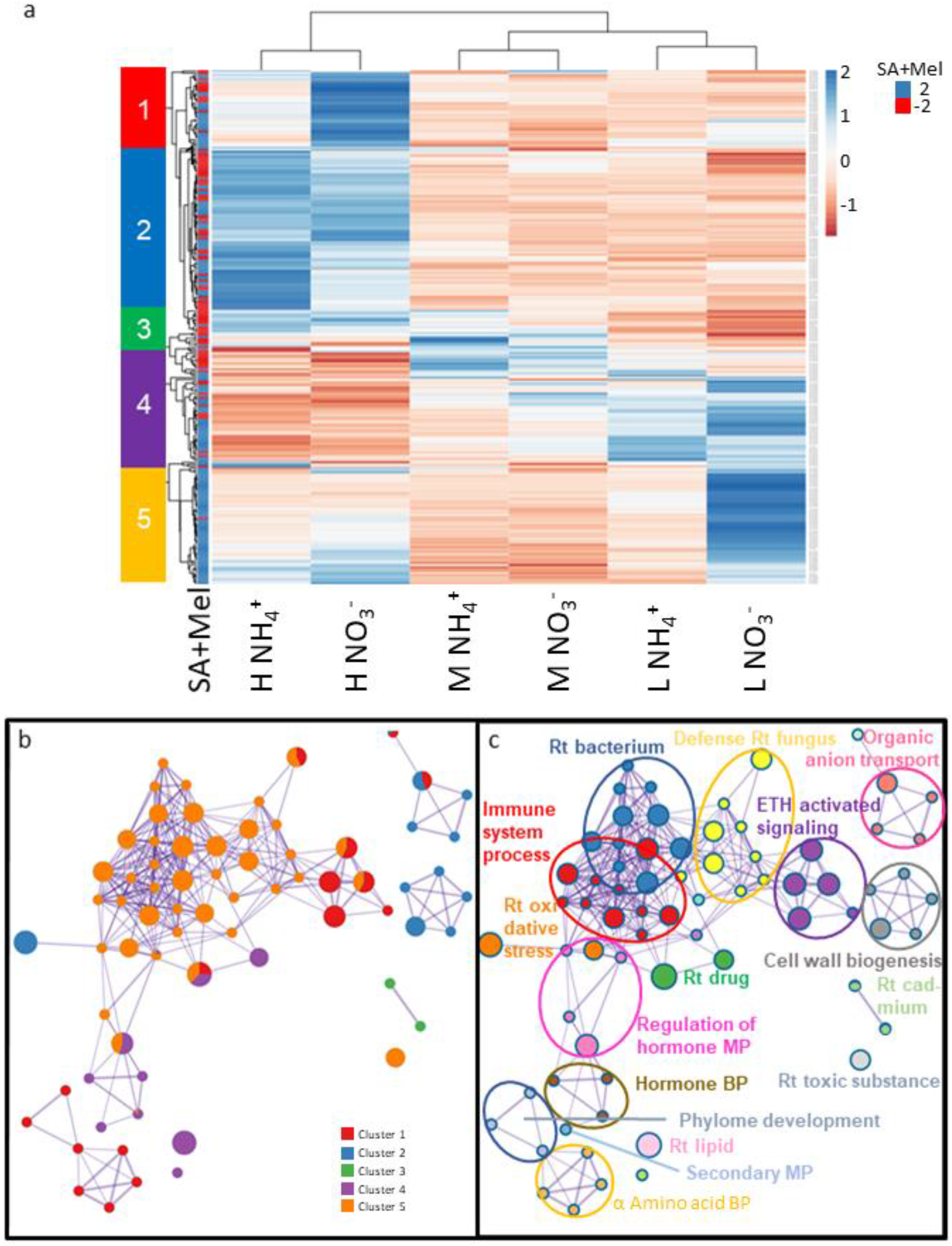
Cluster (a) and network analyses (b,c) of a set of shared differentially expressed genes (DEGs) in leaves of *Populus* x *canescens* exposed to different nitrogen and DEGs responding to application of salicylic acid (SA) and infection with rust fungi (*Melampsora* spp. = Mel). DEGs for SA and Mel responses were extracted from published studies (Luo et al., 2019; Miranda et al., 2007; Rinaldi et al., 2007). a) hierarchical cluster analysis visualizing the pattern of the N-responsive DEGs. MeJA+Chry indicate positive (blue) or negative (red) response of DEGs to SA and Mel treatments. The stacked bar on the left-hand side shows a color code for the five main clusters, b) enrichment network of DEGs within clusters of same color, c) biological processes assigned to DEGs shown in b. Leaves for transcriptome analyses were collected three weeks after feeding poplar with Low (0.4 mM), Medium (2.0 mM) or High (8.0 mM) ammonium or nitrate in the nutrient solution. Bioinformatic analysis were conducted with the best AGI match to the poplar IDs and the standard settings in Metascape. BP = biological process, CP = catabolic process, ETH = ethylene, MP = metabolic process, Rt = response to, n=4 to 5 per treatment.

## Discussion

### Xylem saps shows distinct responses to nitrate and ammonium nutrition

Here, we present the first multi-omics study connecting N nutrition, xylem sap composition and leaf responses. An important result was that changes in N nutrition resulted in profound changes in xylem sap composition. These changes affected not only mobile transport forms of N (nitrate, ammonium, amino acids) and the proteome but all compound classes studied here, including phytohormones, soluble carbohydrates, organic acids, flavonoids, phenolics, benzoates, and many other metabolites. To our very best knowledge the present study reports the largest metabolite library to date available for plant xylem sap and provides in addition to other comprehensive poplar metabolome studies (e.g. Kaling *et al*., 2018; Tschaplinski *et al.*, 2019) a versatile database for future research.

In agreement with previous studies (Siebrecht and Tischner, 1999), we found that a switch from nitrate to ammonium nutrition caused a switch in the dominance of these compounds in the xylem sap. The nitrate concentrations in the xylem sap (about 3 mM) found in our study under low nitrate supply were similar to those reported by Siebrecht *et al.* (2003). In addition to inorganic N, amino acids play a major role as a transport form of N in the xylem sap (Grassi *et al.*, 2002). In poplars under field and laboratory conditions, glutamine is generally the most abundant xylem sap amino-N form (Dickson *et al.*, 1985; Escher *et al.*, 2004; Siebrecht and Tischner, 1999). In our study, glutamic acid-based amino acids dominated as well, but the fractions of glutamine and glutamate in xylem sap could not be distinguished because of cyclization of glutamate and glutamine to pyroglutamic acid (Purwaha *et al.*, 2014) that happened during derivatization prior to GC-MS analysis applied here (Supporting Figure S2).

In soil, the concentrations of nitrate and ammonium vary strongly temporally and spatially, for example, due to varying microbial activities and high mobility of nitrate, thus, requiring flexible adjustment of plant uptake (Glass *et al.*, 2002). Here, we show that the form of applied N resulted in specific effects in some compound classes. For example, high ammonium supply caused a significant increase in xylem sap carbohydrates, whereas the malate concentrations increased under high nitrate. Malate is an important metabolite with major functions during carbohydrate degradation for energy production and via oxaloacetate in anaplerotic reactions. The metabolite ratio of carbohydrates-to-malate regulates photosynthesis by affecting stomatal conductance (Gago *et al.*, 2016; Lima *et al.*, 2019). In line with this notion, we found that the shift in the malate-to-sugar ratio under high nitrate was associated with a much stronger stimulating effect on photosynthesis than high ammonium, although both treatments resulted in similar whole-plant N concentrations.

Most N-containing compounds found in the xylem sap increased with increasing N supply, regardless of the N source but the effects were generally stronger for ammonium than for nitrate nutrition. A novel result was that this behavior was also found for pipecolic acid. Pipecolic acid is not only a degradation product of lysine (Schütte and Seelig, 1967), but its derivate the phytohormone NHP is discussed to be involved in regulating systemic acquired resistance (SAR) (Hartmann *et al.*, 2018; Návarová *et al.*, 2012). Pipecolic acid is intracellularly converted to NHP, which eventually triggers plant immunity responses (Hartmann *et al.*, 2017). Rekhter *et al.* (2019) showed that conversion of pipecolic acid into NHP is not dependent on SA but establishment of SAR may require positive interplay of NHP and SA (Hartmann and Zeier, 2019). Pipecolic acid and NHP move systemically in plants (Návarová *et al.*, 2012; Y.,-C., Chen *et al.*, 2018; Mohnike *et al.*, 2021) but their presence in xylem sap has not been reported yet. Early findings of Návarová *et al*. (2012) suggested NHP rather than pipecolic acid, as the bioactive component inducing SAR. In that light and the notion that pipecolate derives from lysine degradation, the increased amounts of pipecolic acid in high N-treated plants may serve as an additional N transport compound instead of long-distance signaling, a hypothesis that needs to be further elucidated. NHP and SA were low in xylem saps of nitrate-fed plants, regardless of the applied nitrate concentration, whereas NHP increased with increasing ammonium supply. These differences may have implications for SAR. Various studies have shown that SAR signals are influenced by N nutrition (Sun *et al.*, 2020). Our study suggests an intricate role of xylem sap for the transmission of these signals. However, this suggestion needs to be tested in further studies.

Secondary or specialized metabolites play eminent roles in trees species for defense and lignin production (Polle *et al.*, 2013). However, little information is available for the regulation of these compounds in apoplastic compartments including xylem sap. Here, we show that all classes of the confirmed specialized metabolites decreased in poplar xylem sap with increasing N supply, except coumarins. Trade-off between the production of amino- and phenol-based compounds has often been reported and is thought to reflect an ecological plant strategy for carbon utilization balancing growth and defense (Barker *et al.*, 2019). For example, in poplar leaves, specialized metabolites and especially salicinoids are upregulated with decreased N supply (Bryant *et al.*, 1987; Lindroth and Hwang, 1996). Thereby, the palatability of leaves for a number of herbivores is reduced (Fabisch *et al.*, 2019). However, in xylem sap this clear antagonism in response to changes in N availability has not been reported before. Our bioassays underpinned that changes of xylem sap composition strongly affected the proliferation of the endophytic bacterium *B. salicis*. The growth conditions were most favorable in xylem sap of high ammonium-fed poplars, which contained the highest ammonium, amino acid and carbohydrate levels, while the abundance of phenol-based compounds was low. Gram-negative bacteria, the category to which *B. salicis* is belonging to, prefer ammonia over other N-sources (Reitzer, 1996). *Erwinia amylovora*, a close relative of *B*. *salicis* (former *E*. *salicis*), shows increased growth rates under high asparagine levels (Lewis and Tolbert, 1964). It is therefore possible that increasing N sources in the xylem sap promoted the bacterial growth in our study. Obviously, defense proteins, which were also enhanced in the xylem sap of high ammonium-fed poplars, could not attenuate growth of *B. salicis*. Our results imply that *B. salicis* is adapted to vascular conditions and provide valuable guidance for future mechanistic analysis of factors that regulate the interaction of vascular endophytes with their host environment.

### Nitrogen availability affects signaling and defense compounds in xylem sap

Many compounds present in the xylem sap have functions in signaling and defenses (Carella *et al.*, 2016; Turnbull and Lopez‐Cobollo, 2013). One of the most studied phytohormones in xylem sap is ABA. ABA xylem sap concentrations increase under drought (Hansen and Dörffling, 1999; Korovetska *et al.*, 2014; Schachtman and Goodger, 2008). A study with *Ricinus communis* found that the supply with NH_4_^+^ instead of NO_3_^−^ (1 mM) also caused a pronounced increase of ABA in the xylem sap (Peuke *et al.*, 1998). However, in our study application of ammonium instead of nitrate did not affect ABA levels. Since our plants were well irrigated, we did not expect changes in ABA concentrations in the xylem sap.

Here, other phytohormones such as SA, SAG and JA-Ile showed strong responses to poplar N nutrition. These phytohormones have been studied predominantly in xylem sap of crops. For instance, they occur in xylem sap of *Brassica napu*s in the concentration range from 20 to 200 nM (Ratzinger *et al.*, 2009). In poplar xylem sap, we found slightly higher concentrations of these compounds (30 to 600 nM). JA-Ile, SA and its derivative SAG decreased in xylem sap of plants supplied with high N. Both Ja-Ile and SA play intricate – but partly divergent – roles in signaling of plant biotic stress (Pieterse et al., 2014, Sun et al., 2020). Here, we found drastic reprogramming of immune responses in leaf transcriptomes that overlapped with biotic stress responses. Our results suggest that xylem sap phytohormones might be involved in mediating these responses but functional analyses are necessary to test this suggestion.

The xylem sap proteome is dominated by proteins required to maintain the integrity of the xylem architecture (cell wall-related enzymes, general metabolism) and a wide array of proteins such as peroxidases, proteases, chitinases, etc., required as first line of stress defense against invading pests (Rodríguez-Celma *et al.*, 2016). The protein concentrations in xylem saps are low, ranging from 5 and 12 μg mL^−1^ across various herbaceous and tree species examined so far (Rodríguez-Celma *et al.*, 2016). These concentrations are in line with our findings in poplar xylem sap, with the exception of high ammonium treated plants, where the protein concentrations amounted more than 20 μg mL^−1^. In xylem saps of different plant species altogether, approximately 350 non-redundant proteins were detected (Rodríguez-Celma *et al.*, 2016). Previous studies identified 97 proteins in *P*. *trichocarpa* x *P*. *deltoides* (Dafoe and Constabel, 2009) and 102 proteins in *P. deltoides* xylem sap (Pechanova *et al.*, 2010). With the identification of 289 proteins, we report the largest proteome database in poplar xylem sap, so far. Mapping these sequences on *Arabidopsis* gene loci as done by Rodríguez-Celma et al. (2016), still 222 unique proteins were found. Dafoe and Constable (2009) used a pressure extraction method and reported that 33% of the identified proteins contain a signal peptide. In our study, in which contamination by intracellular proteins was extremely low, we found that 63% of the proteins contained a signal peptide. This fraction is similar to those (57%) reported for *P. deltoides* xylem sap (Pechanova *et al.*, 2010) and for the average of xylem sap proteins across different plant species (Rodríguez-Celma *et al.*, 2016). The functional composition found here was also similar to that reported before (Carella *et al.*, 2016; Rodríguez-Celma *et al.*, 2016), supporting species-independent conserved functions of xylem sap proteomes. Our finding that higher N supply enriched the xylem sap with proteins of similar functional classes additionally points to conserved xylem sap functions. However, under high N a number of xylem sap proteins showed significant treatment effects, e.g., differential accumulation of peroxidases, chitinases, and dirigent proteins. Proteins in these classes also accumulated in the xylem sap of crop plants upon attack by a vascular pathogen (Floerl *et al.*, 2012; Yang *et al.*, 2020). The N-dependent regulation of the abundances of xylem sap proteins with anti-microbial characteristics and the enhancement of SAR-related phytohormones suggest local (xylem sap) and systemic (leaves) changes of poplar pathogen resistance.

### Plant-available nitrogen modules systemic defense responses in leaves

Our study highlights that xylem sap is not only a transport medium for water and nutrients but a complex matrix containing multiple signaling and defense compounds that show drastic responses to changes in nitrogen nutrition. There is growing awareness that nitrogen, especially nitrate, modulates plant defenses against pathogens (Mur *et al.*, 2017; Sun *et al.*, 2020; Xuan *et al.*, 2017), in addition to its role as a nutrient (Euring et al., 2014; Luo et al., 2015; Wei et al., 2013, Lu et al. 2019, Gan et al. 2016). Here, we demonstrate that low and high N concentrations systemically regulated poplar leaf defense responses, recruiting JA- and herbivory-related as well as SAR- and pathogen-related pathways. SAR and pathogen-related defenses invoke SA and NHP signaling (Hartmann and Zeier, 2019). These phytohormones increased in xylem sap with increasing nitrogen supply, especially in case of NH_4_^+^. If these xylem sap-transported phytohormones were directly invoked in transcriptional defense activation in leaves, we would have expected GO terms such “Immune system response”, or “Response to fungus”. However, this response was not found under high N, but instead in leaves of plants grown with low nitrate, which contained only moderate SA or NHP levels in the xylem sap. Therefore, direct defense activation by xylem sap SA or NHP is unlikely and the mechanism how xylem sap composition of low nitrate-supplied plants mediates transcriptional activation of defense genes against biotrophic pathogens remained elusive.

Notably, under low nitrate availability also a significant number of genes associated with the GO term “JA-mediating signaling pathway” were differentially expressed in leaves. These genes were also positively regulated in response to poplar leaf beetle feeding and MeJA exposure (Kaling *et al.*, 2018; Luo *et al.*, 2019) and corresponded here to enhanced levels of JA-Ile. The presence of JA-Ile in xylem sap was unexpected because previously only MeJA was identified in xylem sap (tomato, *Thorpe et al.*, 2007). MeJA is a volatile and phloem-mobile long distance signal (Heil and Ton, 2008), while JA-Ile is the bio-active compound for intracellular signaling (e.g. Haroth *et al*., 2019; Thurow *et al*., 2020). Based on the present observation, it possible that JA-Ile might also serve as a systemic signal in the xylem sap.

In conclusion, we demonstrate the potential of xylem sap to transmit nutritional and hormonal signals that result in differential defense activation depending on the nutritional status. Our study does not suggest trade-off between biotrophic and necrotrophic/herbivore defenses because both strategies were simultaneously transcriptionally activated under low N, especially under low NO_3_^−^. It is unknown which changes in xylem sap composition transmitted these responses. However, our encompassing characterization of the xylem sap identified N-driven differences in the abundance of defense proteins, phytohormones, salicinoid-derived compounds as well as many other specialized and primary metabolites, which may be candidates for reprogramming the leaf transcriptome. The results of our study are ecologically highly relevant because they show activation of various defense pathways under resource limitation, pointing to protection of the photosynthetic organs against a broad spectrum of pests. Our study supports that the xylem sap composition contributes to mediate trade-off between defense and growth (Huot *et al.*, 2014) and provides a comprehensive resource for plant researchers.

## Experimental procedures

### Plant propagation and nitrogen treatment

*P.* x *canescens* (hybrid of *P. tremula* x *alba*, clone INRA 717 1B4, INRA, Nancy, France) plants were propagated and cultivated as described by Müller et al. (2013). Plants were *in vitro* multiplied by micro-cuttings and grown on solid, half strength Murashige & Skoog (MS) medium (Murashige & Skoog, 1962) containing in 1 L deionized H_2_O 2.2 g MS medium (Duchefa Biochemie, Haarlem, Netherlands), 10 g sucrose (Duchefa), 0.5 g 2-(N-morpholino)ethanesulfonic acid (ULTRON grade, Merck KGaA, Darmstadt, Germany), and 5 g Gelrite (Duchefa), pH 5.7. The plantlets were kept for four weeks at 23 °C to 24 °C, 60 to 85 μE m^−2^⋅s^−1^ photosynthetically active radiation (PAR) and a 16h/8 h day/night regime. Rooted plantlets were transferred to the greenhouse and further cultivated at 21 °C under ambient light with additional illumination of 150 μE m^−2^⋅s^−1^ PAR and a 16/8 h day/night cycle. They were potted into 0.65 L pots (Lamprecht-Verpackungen GmbH, Göttingen, Germany) containing a mixture of washed and twice autoclaved sand (0.71 to 1.21 mm and 0.4 to 0.8 mm particle size (8:2 v/v), Dorfner GmbH, Hirschau, Germany) and soil (Fruhstorfer Erde “Nullerde”, Hawita Gruppe GmbH, Vechta, Germany) at a ratio of 10:2 (v/v). For acclimation to greenhouse conditions, the plants were covered for one week individually with a transparent plastic beaker, which was gradually lifted during the second week and removed after 14 days. During the acclimation phase, all plants were daily irrigated with modified Long Ashton (LA) nutrient solution (Supporting Table S4, after Hewitt, 1952) containing 2 mM KNO_3_ as N source. Excess nutrient solution ran off. The acclimated plants were divided into two groups and watered daily with modified Long Ashton nutrient solution, containing either ammonium (2 mM NH_4_Cl) or nitrate (2 mM KNO_3_) as N sources (Supporting Table S4). The irrigation volume (per day) was increased with increasing plant height (>10 cm, 50 mL; 11 to 30 cm, 100 mL; 31-60 cm, 150 mL; >61 cm, 250 mL). After 4 weeks, the plants were transferred into 3 L pots (Lamprecht-Verpackungen GmbH) containing a soil/sand mixture (as above). After eight weeks of ammonium or nitrate pre-treatment, each group was split into three subgroups, which were supplied with either 0.4 mM (L), 2.0 mM (M) or 8.0 mM (H) NH_4_Cl or KNO_3_ for three weeks. Each of the six treatment groups consisted of 10 to 12 individual plants, which were randomized regularly. The experiment was repeated three times but not all measurements were possible in all replications. Therefore, the number of biological replicate per measurement varies as indicated in figure and table legends.

### Phenotypic measurements

The stem was marked 2 cm above ground. Growth measurements started 14 days after cultivation of the plants in the greenhouse and were conducted regularly once a week. Plant height was measured with a folding ruler and stem diameter was measured with an electronic caliper (Tchibo GmbH, Hamburg, Germany) at the marked position. The final measurements were conducted at the day of harvest.

### Gas exchange measurements

Gas exchange was measured one day before harvest. A portable photosynthesis system (LI-6800, LI-COR Biosciences GmbH, Bad Homburg, Germany) was attached to a fully developed top leaf with a blade length longer than 10 cm, which was not shaded by other leaves. Measurements of net photosynthesis, transpiration, and stomatal conductance were conducted under saturating light conditions (PAR of 800 μE m^−2^ s^−1^), at ambient CO_2_ concentrations (387 ± 17 ppm) and a mean leaf temperature of 32.6 ± 4.7 °C. Each leaf was acclimated to the measuring conditions for 60 s and then measured 3 or 4 times in intervals of 60 s. Mean values of the repeated measurements were calculated and used for further analyses.

### Harvest and xylem sap extraction

After 13 weeks of greenhouse cultivation, the plants (n = 10 to 12 per treatment and experiment) were harvested randomly within two days, starting after 3 h in the light phase. The shoot was cut 5 cm above the soil surface. The shoot was immediately separated into bark, wood, and leaves. Aliquots of tissues were weighed (lowest 5 cm of the debarked stem, a fully expanded 8 cm-long leaf), shock-frozen in liquid N and stored at −80°C. Three leaves, one fully expanded from the top, one from the middle section and one from the lower part of the plant were weighed separately, dried and anused for N analyses. All further leaves and the remaining stem were weighed separately, dried in paper bags for 7 days at 60 °C and weighed again. Dry biomass of the plant was determined, taking the mass of the samples removed for other analyses into account.

For xylem sap collection, a modified protocol using the natural root pressure was used (Schurr, 1998). Immediately after removing the shoot, a strip of the top 1.5 cm of bark of the stump was peeled off after cutting through the bark and cambium with a razor blade. The exposed wood was washed with deionized H_2_O and blotted with a paper towel. When the first drops of xylem sap accumulated on the cut wood surface, they were removed by washing with deionized H_2_O and the stump was blotted. Then, a 4 cm long piece of tight-fitting tube (TYGON^®^-Tubing for Medical Engineering, RCT GmbH & Co., Heidelberg, Germany) was slipped over the exposed wood and secured at the stump with sticky tape (Durapore^®^, 3M Deutschland GmbH, Neuss, Germany). The sap was pipetted into a 1.5 ml ice cold reaction vessel (Eppendorf AG, Hamburg, Germany) every 10 minutes using a separate glass pipette for each plant. The xylem sap was collected for exactly 4 h and the samples were stored at −80 °C. After xylem sap collection, the roots were washed, weighed, and dried for biomass determination.

### Assessment of xylem sap contamination with intracellular compounds

To test cytosolic contamination, we analyzed glucose-6-phosphate dehydrogenase (EC 1.1.1.49) activities in wood and xylem sap similar as described previously (Polle *et al.*, 1990). Frozen wood was milled (Ball mill MM200, Retsch GmbH, Haan, Germany) whilst cooling with liquid N_2_. Then, 300 mg of the wood powder or 300 μL of the xylem sap were added to 2 mL of extraction buffer (0.1 M sodium phosphate buffer, containing 0.5 % Triton X-100 (Serva Electrophoresis GmbH, Heidelberg, Germany) and 200 mg insoluble polyvinylpyrrolidone, pH 7). The extraction buffer was prepared one day before use to enable swelling of the insoluble polyvinylpyrrolidone and kept at 4°C. Samples were extracted by mixing, incubation for 15 min on ice and centrifugation at 15000 x *g* at 4°C for 30 min. Then 500 μl of the supernatant was filtered through a gel column (NAP 5 Sephadex^®^, Thermo Fisher Scientific Inc., Waltham, Massachusetts, USA), which had been equilibrated and eluted with 1000 μL tris(hydroxymethyl)aminomethane (Tris)-buffer (50 mM Tris HCl (Roth), 1 mM MgCl (Merck KGaA), pH 8.1). Glucose-6-phosphate activity was determined spectrophotometrically (Bergmeyer, 1974). For this purpose, 500 μl Tris-buffer, 300 μl filtered extract, 20 μl 40 mM glucose-6-phosphate (Sigma-Aldrich, now Merck KGaA) and 20 μl 35 mM NADP^+^ (KMF OptiChem, Lohmar, Germany) were added to a cuvette, mixed vigorously and measured at the wavelength of 341 nm and at room temperature. The production of NADH was recorded for 6 min and the increase in absorbance was used to calculate the enzyme activity with an extinction coefficient of 6.22 mM^−1^ cm^−1^.

Glucose-6-phosphate dehydrogenase activities accounted for 0.867 ± 0.065 nkat g^−1^ fresh weight (n = 3) in wood and were not detected in xylem sap (−0.002 ± 0.021 nkat ml^−1^, n = 3). If we assume that glucose-6-phosphate activity was evenly present in the fluid of fresh wood (about 70%), it corresponded to 1.24 nkat ml^−1^. With the maximum activity of glucose-6-phosphate dehydrogenase in xylem sap of 0.02 nkat ml^−1^, the potential contamination of xylem sap with cellular compounds from wood was below 1.6%. We further corroborated by proteome analyses that glucose-6-phosphate was not detected in xylem sap (see below). Therefore, cytosolic contamination of the xylem sap was negligible.

### Carbon and nitrogen analyses

Aliquots of dry leaf, stem, and root tissue were ground to a fine powder in a ball mill (20 s for leaves, 30 s for roots and stems, frequency 30, Type MM400, Retsch, Haan, Germany). Five mg of each sample were weighed into a tin cartouche (IVA Analysentechnik GmbH & Co.KG., Meerbusch, Germany). Carbon and N concentrations were determined with an element analyzer (Vario Micro Cube™, Elementar Analysesysteme GmbH, Langensebold, Germany). Acetanilide (Sigma-Aldrich), was used as the standard.

### NH_4_^+^ and NO_3_^−^ analysis in xylem sap

Xylem sap ammonium and nitrate concentrations were determined with Spectroquant^®^ kits for ammonium or nitrate (Merck KGaA) spectrophotometric analyses according to the instructions of the manufacturer using 100 μl xylem sap. Calibration curves ranging from 0 to 240 nmol N in 100 μl were generated with the Spectroquant kits with NH_4_Cl or KNO_3_ and used to calculate the concentrations of NH_4_^+^ and NO_3_^−^ in xylem sap.

### Measurements of xylem sap protein concentrations

Xylem sap protein concentrations were determined with Pierce^®^ Coomassie Plus kit (Thermo Fisher Scientific), which is based on the Bradford method (Bradford, 1976). Fifty μl xylem sap and 50 μl Coomassie Plus were mixed in a flat bottom 96 well micro titer plate (Greiner AG, Kremsmünster, Austria) and incubated for 10 min at room temperature. During incubation, the plates were centrifuged for 5 min at 4000 x *g* to remove bubbles. Absorbance was measured at 595 nm in a plate reader (Infinite M200 Pro^®^, Tecan Group AG, Männedorf, Switzerland). A dilution series ranging from 1.25 μg ml^−1^ to 10 μg ml^−1^ bovine serum albumin (BSA, Merck KGaA) was used for calibration and processed together with the samples.

To test, if the protein assays were disturbed by low molecular weight compounds in the xylem sap, 0.5 ml aliquots of xylem sap were filtered through Sephadex columns (PD SpinTrap^®^ G-25, GE Healthcare, Chicago, Il, USA) using Tris-buffer (50 mM Tris HCl (Roth), 1 mM MgCl (Merck KGaA), pH 8.1) for column equilibration and sample elution. The protein concentrations were measured as above in the filtered extracts. The mean protein concentration differences between filtered and non-filtered xylem sap samples were less than 20% and not significant. Therefore, we used xylem sap without pretreatment for protein determination.

### Xylem sap metabolites and metabolomics

#### Targeted primary metabolite analysis

Primary metabolites were analyzed by a targeted gas chromatography – mass spectrometry (GC-MS) method. Chemicals and H_2_O used for sample processing were analytical grade (Fisher Scientific). The xylem sap samples were prepared for GC-MS as described by Rotsch et al. (2017). For this purpose, 50 μl unprocessed xylem sap were transferred to a 2 mL reaction tube (Eppendorf) and 200 μL extraction medium (methanol : chloroform : H_2_O, 32.75 : 12.5 : 6.25, v/v/v) and 50 μL *allo*-inositol (0.05 mg μl^−1^, diluted 1:5 in H_2_O) were added. After mixing and centrifugation for 2 min at 20.000 x *g*, 80 μL of the upper phase were transferred to a new 2 mL reaction tube (Eppendorf) and evaporated under constant N_2_ stream. For metabolite derivatization, 15 μL MOX (30 mg methoxamine in 1 mL pyridine) solution was added and the samples were incubated overnight, in the dark and at room temperature. For trimethylsilylation of functional groups, 30 μL N-methyl-N-(trimethylsilyl)trifluoroacetamide were added and the sample was incubated for 1-6 h. Sample separation and fragmentation was done according to Hofvander et al. (2016). For this purpose, 1 μL of sample was injected into the GC (Sigma-Aldrich 5973 Network mass selective detector attached to an Agilent 6890 gas chromatograph (Agilent Technologies, Santa Clara, CA, USA), equipped with a capillary HP5-MS column (Agilent Technologies, J&W Scientific, Folsom, CA, USA). Using internal standards, the Golm metabolome database (GMD) (Kopka *et al.*, 2005) and the National Institute of Standards and Technology (NIST) spectral library 2.0f (NIST, 2005, Gaithersburg, MD, USA), metabolites were identified with the software MSD ChemStation D.01.02.16 (Agilent Technologies). Relative quantification was done by ion count intensity comparison to the internal standard *allo*-inositol.

### Non-targeted metabolite fingerprinting

The metabolites of the xylem sap were extracted as described by Feussner & Feussner (2019) in the chapter ‘Extraction of metabolites from aqueous solutions of plants and fungal origin’. Non-targeted metabolite fingerprinting was conducted by liquid chromatography coupled to high resolution-mass spectrometry (LC-HRMS) as described in detail by Feussner & Feussner (2019). Two independent experiments were performed for non-targeted metabolome analysis of xylem sap. One experiment was analyzed by ultra-high-performance LC (UHPLC, LC 1290 Infinity (Agilent Technologies) coupled to a HRMS (6540 UHD Accurate-Mass Q-TOF LC-MS instrument (Agilent Technologies). The second experiment was analysed by UPLC (Waters Corporation) coupled to TOF-MS LCT Premier (Waters Corporation). The settings applied for the analyses of the samples were described in detail by Feussner and Feussner (2019). Data acquisition of accurate mass data was performed either with Mass Hunter Acquisition B.03.01 (Agilent Technologies) and analyzed with Mass Hunter Qualitative Analysis (Agilent Technologies) or with MassLynx 4.1 (Waters Corporation). Peak picking and alignment were performed with Profinder B.08.00 (Agilent Technologies) or with MarkerLynx (Waters Corporation) software. The metabolite features covered in both experiments were significantly correlated (Supporting Figure S5). Metabolite annotation was done by database search based on the accurate mass information. The chemical identity of compounds were determined by interpretation of HRMS/MS spectra (Abreu et al., 2020; Green et al., 2020). For further data processing, statistics, visualization as well as for data mining the MarVis-Suite toolbox (http://marvis.gobics.de/, Kaever et al., 2015) was used. The raw data were deposited in the Metabolights database (https://www.ebi.ac.uk/metabolights/) under the accession number MTBLS2305.

### Phytohormone analyses in xylem sap

An LC-MS approach was used for phytohormone analysis, as described by Herrfurth and Feussner (2020) and Mohnike et al. (2021). For phytohormones extraction, 200 μL unprocessed xylem sap and 750 μL phytohormone solution (10 ng D_4_ JA-Leu (kindly provided by Otto Miersch, Halle/Saale, Germany), 10 ng D_6_-ABA, 10 ng D_4_-SA (both from C/D/N Isotopes Inc., Pointe-Claire, Canada), 50 ng D9-Pip (Merck KGaA) and 50 ng ^13^C_6_-SAG (kindly provided by Prof. Petr Karlovsky, Goettingen, Germany) in 750 μL methanol) were mixed. The phytohormones were extracted as described for non-targeted metabolite fingerprinting (as described above). The combined upper phases were evaporated under constant N_2_ stream and re-suspended in 100 μL of acetonitrile : H_2_O (20 : 80, v/v), containing 0.3 mM NH_4_HCOO (pH 3.5, adjusted with formic acid). An ultra-performance liquid chromatography system (ACQUITY UPLC™ system, Waters Corp.) was used for reversed phase separation, followed by nano-electrospray ionization (nanoESI) with a chip ion source (TriVersa Nanomate^®^; Advion BioSciences, Ithaca, NY, USA). Ionized phytohormones were determined with an AB Sciex 4000 QTRAP^®^ tandem mass spectrometer (AB Sciex, Framingham, MA, USA) in negative and positive mode, respectively, using mass transitions and optimized parameters shown in Supporting Table S5. Phytohormone quantification was performed using an internal standard based calibration curve including the data intensity (*m*/*z*) ratios of unlabeled/deuterium-labeled *versus* molar amounts of unlabeled standards.

### Proteomics

All solvents used for the processing of xylem sap and the resulting proteins were liquid chromatography-mass spectrometry (LC-MS) or high-performance liquid chromatography (HPLC) grade and obtained from Fisher Scientific (Hampton, NH, USA). 400 μL xylem sap were lyophilized (Freeze dryer, Piatkowski Forschungsgeräte-Vertrieb, München, Germany) for 5 days. For protein denaturation and to reduce disulfide bonds, the samples were dissolved in 60 μL ammonium hydrogen carbonate (ABC, Fluka Analytical, now Sigma-Aldrich and further-on called Sigma-Aldrich) containing D-1,4-dithiothreitol (DTT, Sigma-Aldrich) (100 mM ABC, 100 mM DTT, 1:4, v/v) and 75 μL 2,2,2-trifluoroethanol (TFE) and incubated for 30 min at 60 °C in a water bath. Samples were centrifuged at 16.000 x *g* for 10 min and 100 μL supernatant were transferred to a 1.5 mL low binding micro reaction tube (Eppendorf). For SH-group alkalization, 5 μL 500 mM iodoacetamide (IAA, Sigma-Aldrich) were added and the sample was incubated in darkness for 30 min. Subsequently, 50 μL H_2_O were added to dilute the sample. For protein precipitation, 620 μL methanol and 200 μL chloroform were added and the sample was carefully mixed after each addition (Wessel and Flügge, 1984). Subsequently, 70 μL 625 mM NaCl (Merck KGaA) and 395 μL H_2_O were added. The sample was mixed and centrifuged for 20 min at 16100 x *g*. The proteins accumulated in the interphase. The upper phase was removed by pipetting without disturbing the inter phase. Five hundred μL methanol were added, the sample was mixed and centrifuged for 20 min at 16.000 x *g*. Thereby, a protein pellet was formed. The supernatant was carefully decanted. The protein pellet was dried at room temperature for about 25 min and then dissolved in 50 μL 100 mM Tris HCl, pH 8.0 (Roth) by sonication for 3 s. For protein digestion, 1 μL trypsin solution (100 ng μL^−1^ Sequencing grade, Roche Deutschland Holding GmbH, Mannheim, Germany) in 10 mM HCl (Merck KGaA) was added and the sample incubated for 16 h at 37 °C in a water bath. Digestion was stopped by addition of 20 μL 20 mM ammonium formate (AF, Sigma-Aldrich). The sample was centrifuged for 20 min at 16.000 x *g* and the peptides in the supernatant were purified.

For peptide purification, Stop and go extraction (STAGE) tips (Rappsilber *et al.*, 2003) were used. A triple layer of the C18 matrix (C18 Empore™ Disks, 3M) was inserted into 20 μL pipette tips (Gilson Company Inc., Columbus, OH, USA) and this STAGE tip mounted on a 0.5 μL micro reaction tube (Eppendorf) in a centrifuge (5415R, Eppendorf). The STAGE tip was loaded with solvents by adding subsequently the following liquids, each followed by centrifugation: 10 μL methanol, 1 min at 1.800 x *g*, 20 μL acetonitrile, 1 min at 1.800 x *g*, and 20 μL AF, 4 min at 1.800 x *g*. Subsequently, 60 μL of peptide solution were added and centrifuged for 10 min at 1.800 x *g*. 15 μL 20 mM AF were added and centrifuged for 2 min at 1.800 x *g*, followed by 2 min at 2.600 x *g*. For peptide elution, the STAGE tip was placed in a new, low-bind 0.5 μL micro reaction tube (Eppendorf) and 40 μL of 60% acetonitrile in 20 mM AF, pH 10 were added to the tip. The STAGE tip was centrifuged for 4 min at 1.800 x *g*, followed by 2 min at 3.000 x *g*. The eluted peptide solution that had passed through the STAGE tip was evaporated in a vacuum concentrator (Eppendorf). Dried peptides were dissolved in 2 % acetonitrile, 0.1 % formic acid in H_2_O in a volume to obtain a final concentration of 6 ng peptides in 100 μL solvent, taking into account a 30% loss during processing.

Further sample processing was performed in the core facility LCMS Protein Analytics of the University of Goettingen according to the following pipeline used in the study of Horianopoulus et al. (2020): “2 μL of each sample were used for liquid chromatography mass spectrometry (LCMS) analysis. Peptides were loaded on an Acclaim PepMap 100 pre-column (100 μm x 2 cm, C18, 3 μm, 100 Å; Thermo Fisher Scientific) with 0.07% trifluoroacetic acid at a flow rate of 20 μL min^−1^ for 3 min. Analytical separation of peptides was performed on an Acclaim PepMap RSLC column (75 μm x 50 cm, C18, 3 μm, 100 Å; Thermo Fisher Scientific) at a flow rate of 300 nL min^−1^. The solvent composition was gradually changed within 94 min, starting from 96% solvent A (0.1% formic acid) and 4% solvent B (80% acetonitrile, 0.1% formic acid) to 10% solvent B within 2 minutes, to 30% solvent B within the next 58 min, to 45% solvent B within the following 22 min, and to 90% solvent B within the last 12 min of the gradient. All solvents were Optima grade for LC-MS (Thermo Fisher Scientific). The eluting peptides were on-line ionized by nano-ESI deploying the Nanospray Flex Ion Source (Thermo Scientific) at 1.5 kV (liquid junction) and transferred into a Q Exactive High Field mass spectrometer (Thermo Fisher Scientific). Full scans in a mass range of 300 to 1650 *m*/*z* were recorded at a resolution of 30,000, followed by data-dependent top 10 higher-energy collisional dissociation fragmentation at a resolution of 15,000 (dynamic exclusion enabled). LC-MS method programming and data acquisition were performed with the XCalibur 4.0 software (Thermo Fisher Scientific).”

The resulting raw data were used for protein identification using MaxQuant Software v1.6.6.0 (Cox & Mann, 2008) with the programs default parameters. Label-free quantification was enabled, trypsin/P was used as digestion mode. For database search and annotation, a JGI/Phytozome12-derived *P. trichocarpa* (v.3.1) specific database and the Andromeda algorithm (MaxQuant v.1.6.6.0) were used. The results of the MaxQuant database search were evaluated using the Perseus software v.1.6.6.0 (Cox and Mann, 2008). Protein functional group assignment was done with MapMan (https://mapman.gabipd.org/, assessed 02/2020, Thimm et al., 2004). Metascape (Zhou *et al.*, 2019) was used for functional group enrichment analysis. The raw data were deposited in the PRIDE database (https://www.ebi.ac.uk/pride/) under the accession number PXD024142.

### Xylem sap *Brennaria salicis* growth assay

A stock culture of *Brennaria salicis* (formerly *Erwinia salicis*) strain PD135 (NCCB 87018) was purchased from the Westerdijk Fungal Biodiversity Institute (Wageningen, Netherlands). *B*. *salicis* was stored in glycerol stock at −80 °C and reactivated on tryptic soy broth (TSB) plates (Art. No. 100550, Merck KGaA). After 24 h incubation at 22 °C in the dark, one inoculation loop of bacterial cells was added to 30 mL liquid TSB. The solution was incubated at 28 °C and 160°rpm for 24°h (Maes et al., 2009). Of the bacterial suspension, 1.5 mL were transferred to a 2 mL reaction vessel (Sarstedt). The bacteria were spun down at 3930 x g (Rotanta 96R, Hettich GmbH & Co. KG, Tuttlingen, Germany) and the supernatant was discarded. Optical density (OD), measured at 600 nm, was adjusted to OD = 1 by diluting the culture with 1 mM MgCl (R. L. Smith & Maguire, 1998). Eight replicates of 100 μl xylem sap of each nitrogen treatment and 8 controls (100 μl water) were added to a flat bottom, 96-well plate (Greiner) and 2.5 μL bacterial solution was added. The plate was introduced to an automated plate reader (Infinite 200Pro, Tecan) and incubated for 24 h using a manufactures protocol (Tecan Trading AG, 2012) with following modification: during incubation, the temperature was set to 28 °C and orbital amplitude to 5 mm. OD was measured at 600 nm every 30 min.

### Leaf transcriptome analysis

An expanded, fully light exposed top leaf that was formed during the N treatments was snap frozen in liquid N_2_ and stored at −80 °C. Leaves from five (8 mM N treatments) or four plants of each treatment were used for RNA extraction. Leaves were milled under liquid N_2_ in a ball mill (MM400, Retsch). The frozen leaf powder of the high N treatments was extracted with the innuPrep Plant RNA kit (Analytik Jena, Jena, Germany), that of the other treatments using the cetyltrimethyl ammonium bromide method (Chang et al., 1993) with slight modifications reported previously (Janz *et al.*, 2012). RNA concentrations were determined spectophometrically (NanoDrop2000, Thermo Fisher Scientific) and ranged from 92 to 2000 ng μL^−1^. We used 10 μL of each sample containing at least 1 μg RNA for RNA sequencing and library generation and 3 μL the determination of RNA integrity numbers (RIN) These analyses were conducted by Chronix Biomedical (Göttingen, Germany), using the RNA Nano Chip Kit (Bioanalyzer 2100, Agilent Technologies) and four samples per treatment with highest RINs were used for further processing (Supporting Table S6). Enrichment of poly-A RNA was performed with the New England Biology (NEB)-Next Poly (A) mRNA Magnetic Isolation Module (New England Biolabs, Ipswich, MA, USA) following the manufacturers’ manual. Seventy-five bp single-read sequences were generated using the NextSeq500 (Illumina, Inc., San Diego, C, USA) instrument. The sequences were used to create libraries with NEBNext Ultra RNA Library Prep Kit for Illumina (New England Biolabs).

### Bioinformatic analysis of RNAseq data

Adapter trimming and quality filtering were done with fastP, using default parameters (S., Chen *et al.*, 2018). Criteria for quality filtering included base accuracy, N base number and read length. This procedure resulted in 13 x10^6^ to 21 x10^6^ reads per sample (Supporting Table S6). Reads were mapped against the *P. tremula* x *alba* 717-1B4 data base (Mader *et al.*, 2016), v. 2, downloaded from http://aspendb.uga.edu) with the read alignment tool Bowtie2 (Langmead and Salzberg, 2012). The relative number of reads successfully mapped to a gene model ranged from 89% to 94% (with one outlier, M NH_4_^+^_2 = 77.47%) (Supporting Table S6). The raw data were deposited in the ArrayExpress database (http://www.ebi.ac.uk/arrayexpress) under the accession number E-MTAB-8930.

Linux command-line utility grep (Torvalds, 2015) was used to extract information from mapping files, which were integrated to count tables with the statistical software R base version (R Studio v. 1.2.5033, R Core Team, 2014). The package DESeq2 (Love *et al.*, 2014) was used to identify differentially expressed genes (DEGs) between the treatments. DEGs per treatment were mapped on KEGG pathways (https://www.genome.jp/kegg/, assessed 02/2020, Kanehisa & Goto, 2000). Gene ontology enrichment analyses were executed with Metascape (https://metascape.org, accessed 06/2020, Zhou et al., 2019) using the Arabidopsis best hit AGIs of the poplar DEGs. To identify genes in our data set with roles in poplar fungal, leaf feeding, and phytohormone interactions, DEGs (log2 > 1, p_adjusted_ < 0.05) published by Kaling et al. (2018), Luo et al. (2019), Miranda et al. (2007) and Rinaldi et al. (2007) were downloaded. The DEGs were filtered and those with consistent up- or down-regulation in response to Me-JA (Luo et al. 2019) and leaf beetle treatment (Kaling et al. 2018) were kept and used to extract shared DEGs of the N treatments. Heatmaps of the shared DEGs were generated with ClustVis (https://biit.cs.ut.ee/clustvis/, Metsalu & Vilo, 2015). Gene ontology (GO) term enrichment analyses and network analyses of the shared DEGs were conducted with Metascape (https://metascape.org, accessed 06/2020, Zhou et al., 2019) using the Arabidopsis best hit AGIs of the poplar DEGs. The same strategy was used to find shared DEGs between N treatments and SA and Melampsora exposure using DEGs published by Luo et al. (2019), Miranda et al. (2007) and Rinaldi et al. (2007). The shared DEGS used in our analyses are presented in Supportive data file S5.

### Statistical analysis

Data are shown as box plots or means (± SD), if not indicated otherwise. The number of biological replicates is indicated in the tables and figures legends. To compare means, we tested normal distribution of the data sets by visual inspection of residuals or used the Shapiro Wilk test when the visual inspection was unclear (Kim, 2013). If data were not normally distributed, they were log transformed to achieve normal distribution. Statistical tests were conducted with R (R Core Team, 2014), using generalized mixed models, Tukey’s test or Welch’s t-test in the package Multcomp (Hothorn *et al.*, 2008). Differences between treatments were considered to be significant, when the p-values of the post hoc test were p < 0.05. The packages ggfortify (Tang *et al.*, 2016) and cluster (Maechler, 2019) were used for principal component analyses. Venn diagrams were generated with InteractiVenn (Heberle et al., 2015, http://www.interactivenn.net/).

## Supporting information

Supportive Figures and Tables

Supportive data file 1

Supportive data file 2

Supportive data file 3

Supportive data file 4

Supportive data file 5

## Acknowledgements

We gratefully acknowledge the excellent technical assistance of Merle Fastenrath, Cathrin Leibecke, Julian Wellhäuser and Mónica Daniela Rodriguez Nava (Forest Botany and Tree Physiology) for supporting plant culture, Sabine Freitag (Service Unit for Metabolomics and Lipidomics) for sample preparation for metabolome analyses and Thomas Klein (Laboratory for Radioisotopes) for RNA extraction. This research was funded by the Deutsche Forschungsgemeinschaft (DFG) (IRTG 2172: PRoTECT, project M2). Proteome analyses was conducted at Service Unit LCMS Protein Analytics of the Göttingen Center for Molecular Biosciences (GZMB) at the Georg-August-University Göttingen (Grant DFG-GZ: INST 186/1230-1 FUGG to Stefanie Pöggeler).

## AUTHOR CONTRIBUTIONS

KK and AP conceived and planned the study. KK performed the experiments, collected and prepared the samples and contributed to the analyses. INA and KF conducted untargeted metabolome analyses and identified metabolites. KZ and CH performed phytohormone analyses. IF supervised the metabolome and phytohormone analyses. TI conducted GCMS analyses of primary metabolites. AM, OV, and KS performed proteome analyses under supervision of GHB. DJ and AP conducted transcriptome analyses. KK synthesized the results, performed statistical analyses and wrote the first draft of the paper. All authors contributed to writing and agreed on the final version of this manuscript.

## Data availability statement

Leaf transcriptome: ArrayExpress database, accession number E-MTAB-8930

Xylem sap metabolome: Metabolights database, accession number MTBLS2305

Xylem sap proteome: PRIDE database, accession number PXD024142

## Supporting Information

Additional Supporting Information may be found in the online version of this article.

| File name | Description |
| --- | --- |
| **Supportive tables**<br>Word document, 56 KB | **Table S1.** Plant dry mass, tissue N concentrations and net photosynthesis of poplars grown with increasing concentrations of ammonium or nitrate<br>**Table S2.** GC-MS detected xylem sap metabolites of *Populus x canescens*, supplied with increasing concentrations of ammonium or nitrate |
|  | **Table S3.** Metabolite profile of *Populus x canescens* xylem sap, identified by targeted (GC-MS, LC-MS) and non-targeted approaches (LC Qtof MS)<br>**Table S4.** Composition of the nutrient solutions used to irrigate *Populus x canescens* during the experiment<br>**Table S5.** Mass transitions and optimized parameters for detection of phytohormones by mass spectrometry<br>**Table S6:** Quality and processing details of RNA sequencing and library generation |
| **Supportive figures**<br>Word document, 380 KB | **Figure S1.** Growth and height of poplars supplied with increasing concentrations of ammonium or nitrate<br>**Figure S2.** Spectra obtained by measuring standards of glutamine and glutamate by GC-MS<br>**Figure S3.** Correlation analyses of ion count intensities of identified metabolites in untargeted approaches of two independent experiments<br>**Figure S4.** Cluster analysis of 863 features obtained by untargeted LC Qtof-MS analysis in xylem sap of *Populus x canescens*, supplied with increasing concentrations of ammonium or nitrate<br>**Figure S5.** Differentially expressed genes (DEGs) in leaf transcriptome data of *Populus x canescens* plants, supplied with increasing concentrations of ammonium or nitrate. |
| **Supportive data file S1**<br>MS Excel, 963 KB | Metabolite features in *Populus x canescens* xylem sap by LC Qtof MS |
| **Supportive data file S2**<br>MS Excel, 21 KB | Detection details of identified *Populus x canescens* xylem sap secondary metabolites by LC Qtof MS and HRMS/MS fragmentation analyses |
| **Supportive data file S3**<br>MS Excel, 271 KB | *Populus x canescens* xylem sap proteome |
| **Supportive data file S4**<br>MS Excel, 58 KB | Differentially expressed genes in leaves of *Populus x canescens* treated with same concentration of either ammonium or nitrate |
| **Supportive data file S5**<br>MS Excel, 923 KB | Comparison of 2-fold up or down DEGs in this study with published studies in response to methyl jasmonate treatment, salicylic acid, a biotrophic pathogenic fungus or herbivory |

