## Supplementary material for "Multi-omics analysis of xylem sap uncovers dynamic modulation of poplar defenses by ammonium and nitrate": Supportive Figures and Tables

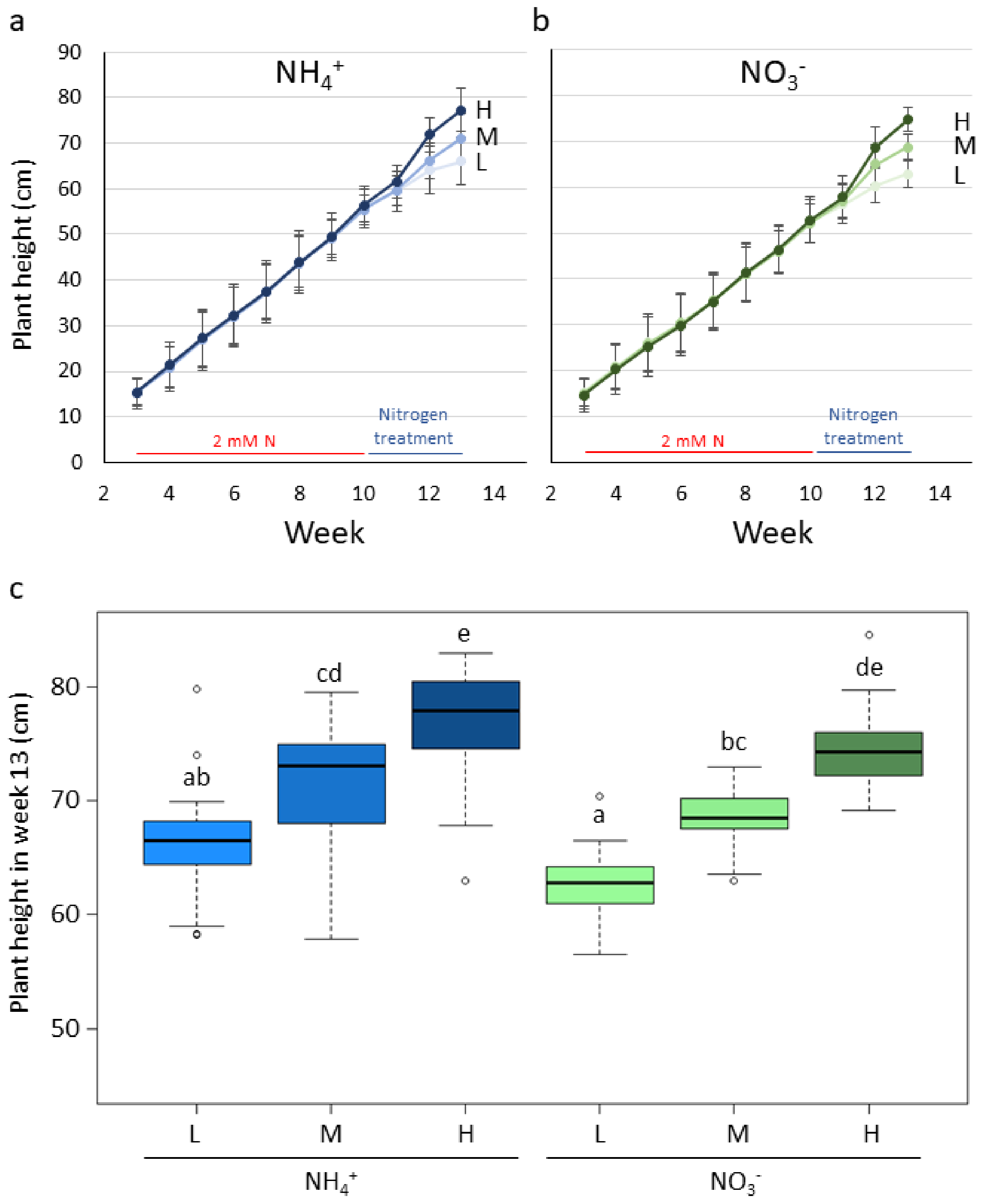


Figure S1: Growth (a,b) and height (c) of poplars supplied with increasing concentrations of ammonium or nitrate. *Populus* x *canescens* plants were grown with 2 mM $NH_4^+$ or $NO_3^-$ for 10 weeks and then supplied for 3 weeks with 0.4 mM (L), 2.0 mM (M) or 8.0 mM (H) $NH_4^+$ or $NO_3^-$ in the nutrient solution. a) Height growth of poplars treated with $NH_4^+$, b) Height growth of poplars treated with $NO_3^-$, c) Height of poplars at the harvest. Different letters above the boxplots indicate significant differences at $p < 0.05$ determined by a generalized mixed model with the biological replicates as random factor. N = 11 per treatment.

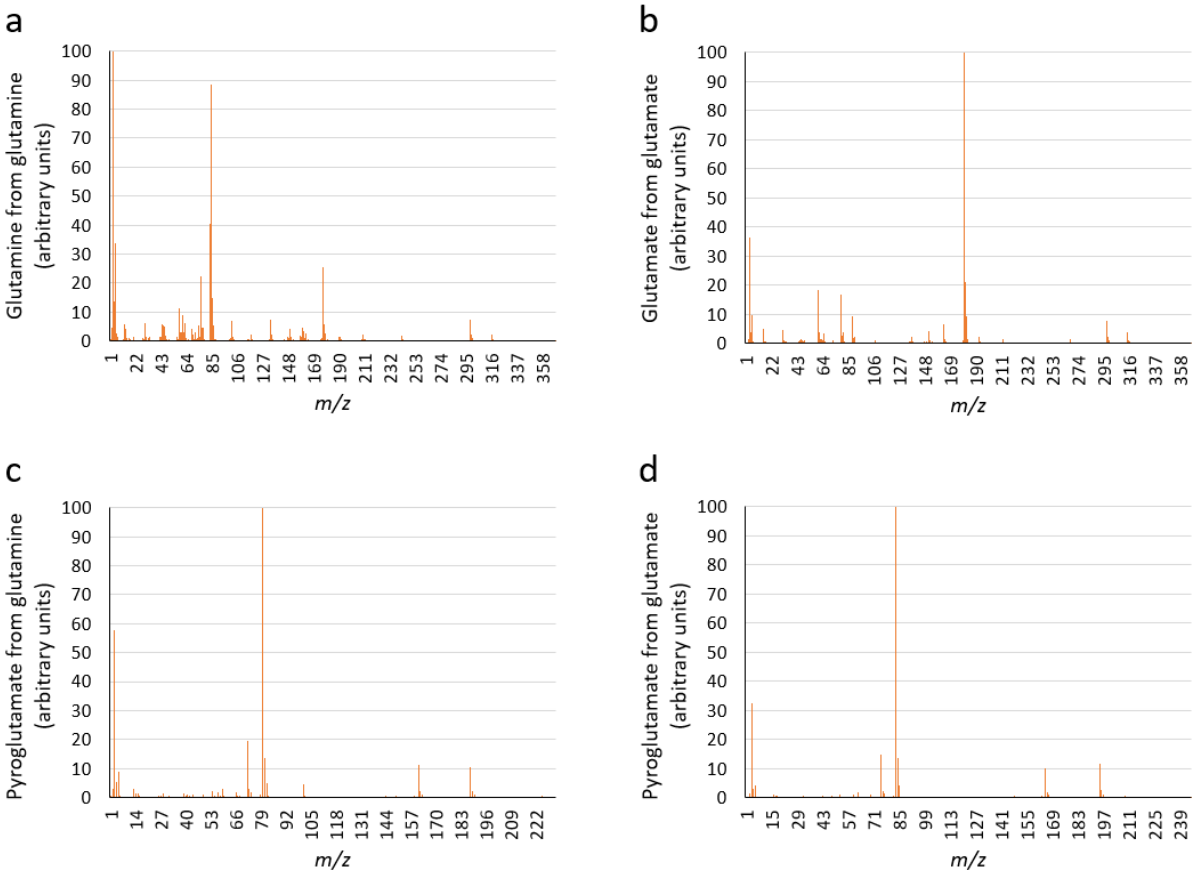


**Figure S2:** Spectra obtained by measuring standards of glutamine and glutamate by GC-MS. Both glutamine and glutamate can be converted to pyroglutamate during derivatization. a) Glutamine (a) and glutamate (b) standards were subjected to the same derivatization procedure applied to analyzed samples. In the GC-MS analysis in both cases not only the three time trimethylsilylated (3 TMS) derivatives were detected but also pyroglutamate (c, d) (2TMS). Displayed are the recorded spectra of the substances that matched the spectra in the NIST library. The highest arbitrary unit value was set to 100 in each diagram.

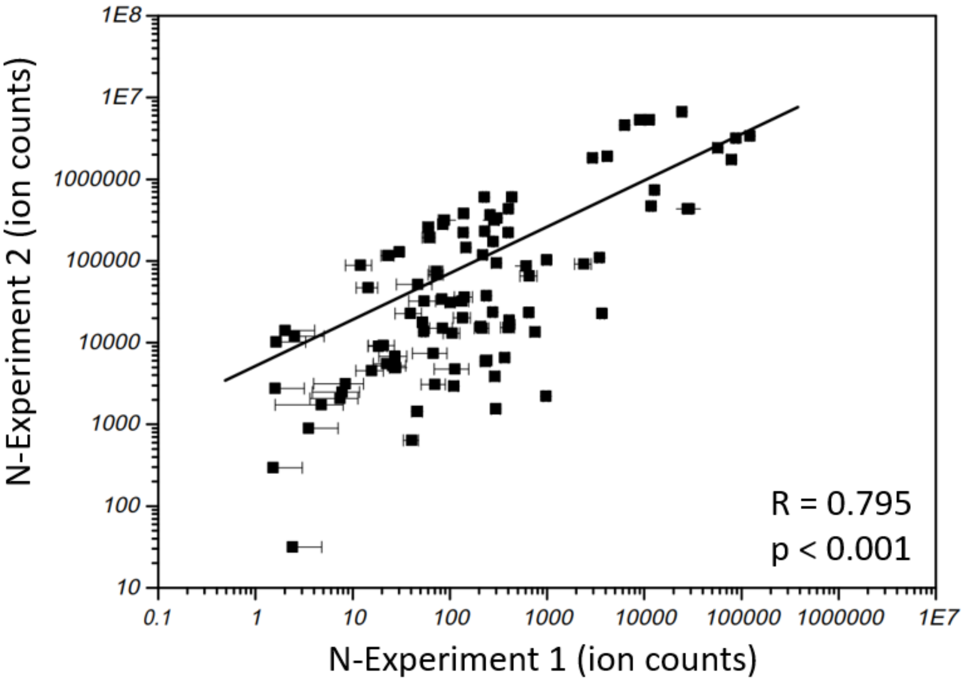


**Figure S3:** Correlation analyses of ion count intensities of identified metabolites in xylem saps analyzed by untargeted approaches of two independent experiments. Samples were analyzed by UHPLC-HRMS in experiment 1 (n= 8 per treatment) and by UPLC-Qtof MS in experiment 2 (n= 6 per treatment). Data show means per compound and experiment. Shared compounds in both experiments are shown. Xylem saps were analyzed after three weeks of feeding poplar with L (0.4 mM), M (2.0 mM) or H (8.0 mM) ammonium or nitrate in the nutrient solution.

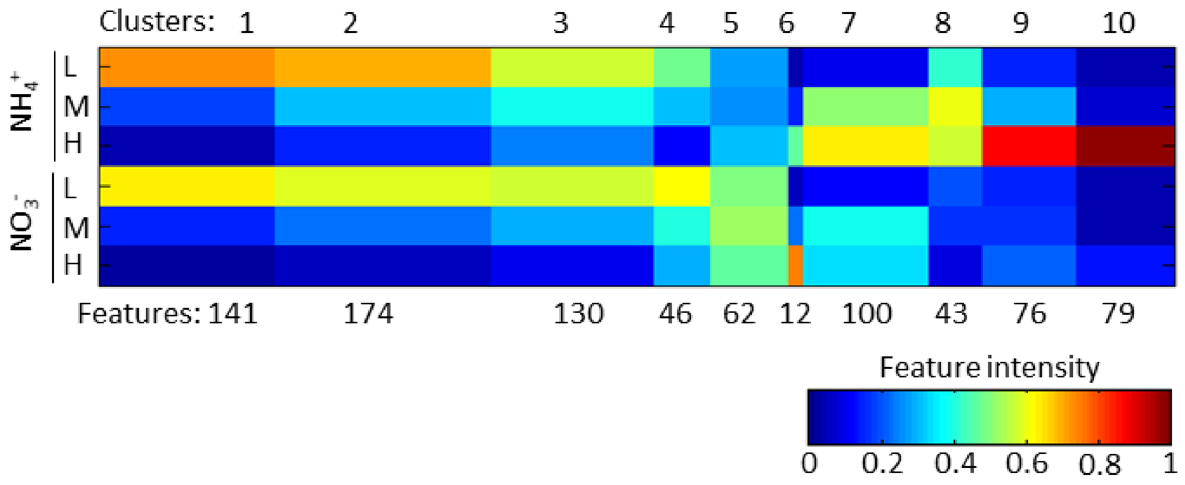


**Figure S4:** Cluster analysis of 863 features obtained by an untargeted metabolomics approach in xylem sap of *Populus* x *canescens*, supplied with increasing concentrations of ammonium or nitrate. The cluster analysis was conducted by MarVis (Kaever et al., 2015). Ten clusters were generated during analysis (numbers above heatmap) containing different numbers of features (numbers below heatmap). Xylem saps were analyzed after three weeks of feeding poplar with L (0.4 mM), M (2.0 mM) or H (8.0 mM) ammonium or nitrate in the nutrient solution. Features that differed significantly between at least two treatments were obtained by ANOVA and subjected to Benjamini-Hochberg correction at $FDR_{adjusted}$ $p < 10^{-6}$. n= 5 to 6 per treatment.

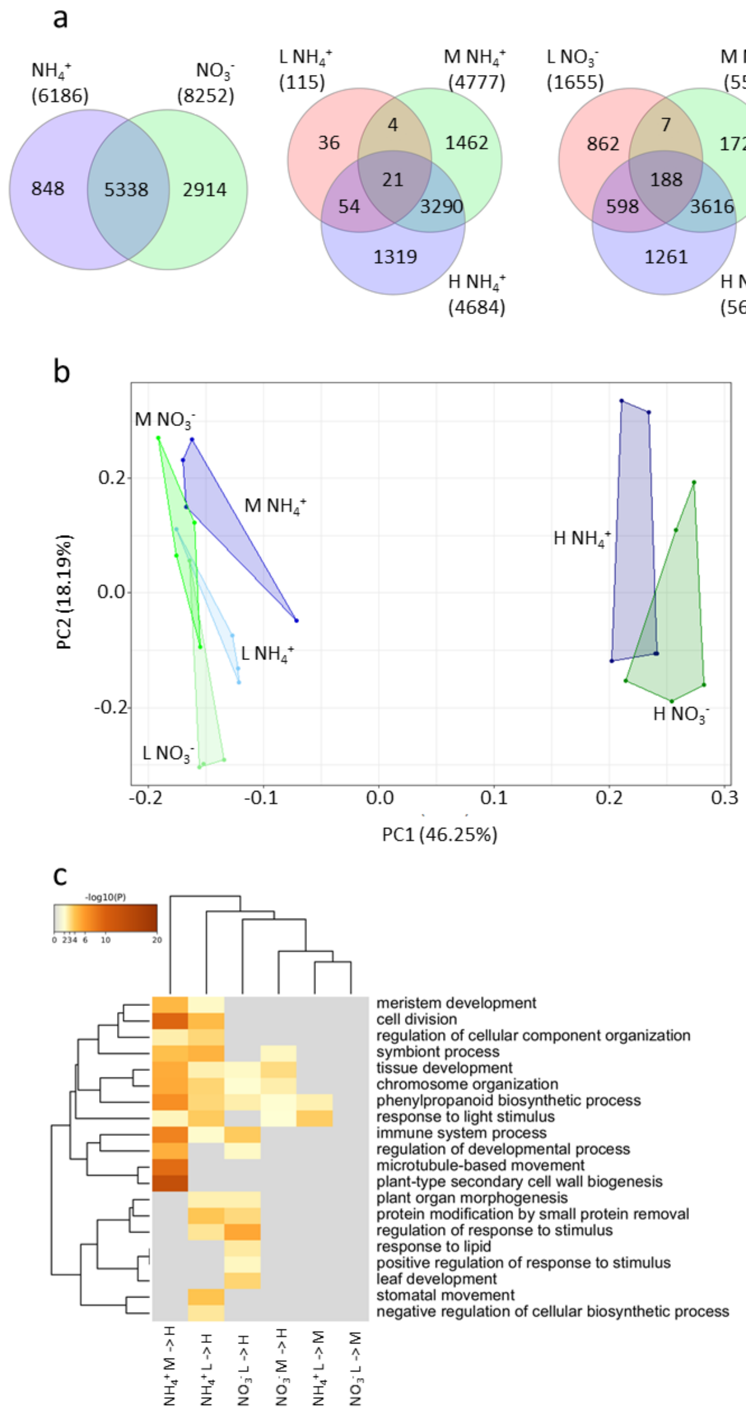


**Figure S5** Differentially expressed genes (DEGs) between at least two treatments in the leaf transcriptomes of *Populus* x *canescens.* Leaves for transcriptome analyses were harvested three weeks of growth with low (L: 0.4 mM), medium (M: 2.0 mM) or high (H: 8.0 mM) ammonium or nitrate in the nutrient solution. a) Venn diagrams showing numbers of unique and overlapping DEGs in leaves of plants supplied with supplied with low, intermediate, or high $NH_4^+$ or $NO_3$, b) Principal component analysis of the DEGs, c) Hierarchical cluster analysis of significantly enriched gene ontology terms in the DEGs. Analyses were conducted with the best AGI matches of the poplar IDs and standard settings in Metascape. Leaves were analyzed after three weeks of feeding poplar with n= 4 to 5 per treatment.

**Table S1**: Plant dry mass, net photosynthesis, and tissue nitrogen concentrations of *Populus* x *canescens* treated with different forms and concentrations of nitrogen

| Treatment | Plant dry mass (g plant$^{-1}$) | | Net PS (µmol m$^{-2}$ s$^{-1}$) | | Root N (mg g$^{-1}$) | | Stem N (mg g$^{-1}$) | | Leaf N (mg g$^{-1}$) | |
|---|---|---|---|---|---|---|---|---|---|---|
| $NH_4^+$ L | 10.73 | ± 1.69 a | 6.26 | ± 2.04 ab | 0.95 | ± 0.07 ab | 0.56 | ± 0.05 a | 1.22 | ± 0.17 a |
| $NH_4^+$ M | 10.78 | ± 1.07 a | 6.45 | ± 2.36 b | 1.09 | ± 0.05 bc | 0.91 | ± 0.02 b | 2.05 | ± 0.31 bc |
| $NH_4^+$ H | 13.27 | ± 1.38 b | 7.95 | ± 3.63 c | 1.62 | ± 0.06 d | 2.07 | ± 0.07 d | 2.5 | ± 0.52 cd |
| $NO_3^-$ L | 10.82 | ± 1.40 a | 4.31 | ± 0.91 a | 0.88 | ± 0.06 a | 0.52 | ± 0.07 a | 1.03 | ± 0.07 a |
| $NO_3^-$ M | 11.24 | ± 1.07 a | 5.69 | ± 1.66 ab | 1.16 | ± 0.07 c | 0.9 | ± 0.05 b | 1.86 | ± 0.18 b |
| $NO_3^-$ H | 13.73 | ± 1.19 c | 12.29 | ± 1.20 d | 1.84 | ± 0.17 e | 1.72 | ± 0.06 c | 2.99 | ± 0.23 d |

$NH_4^+$ = ammonium, $NO_3^-$ = nitrate, PS = photosynthesis, N = Nitrogen. L = 0.4 mM, M = 2.0 mM, H = 8.0 mM nitrogen supplied in nutrient solution. Data show means (± SD) for the following number of replicates per treatment: Net PS n=14, Plant dry mass n=22, Tissue N concentrations n= 4 to 5. Different letters in a column indicate significant differences at $p \leq 0.05$ according to Tukey´s-Kramer post-hoc test or, when data of more than one experimental replicate was analyzed, a generalized mixed model with replicate as random factor.

**Table S2**: GC-MS detected xylem sap metabolites of *Populus* x *canescens*, supplied with increasing concentrations of ammonium or nitrate.

| | | Ammonium | | | Nitrate | | |
|---|---|---|---|---|---|---|---|
| | **Identified by** | **Low** | **Medium** | **High** | **Low** | **Medium** | **High** |
| **Amino acids** | | | | | | | |
| beta-Alanine | ES | 0.81 | 0.89 | 1 | 0.55 | 0.62 | 0.53 |
| 2-Amino butyrate | SL NIST | 0 | 0 | 1 | 0 | 0.01 | 0.42 |
| Alanine | ES | 0.03 | 0.1 | 1 | 0.04 | 0.1 | 0.33 |
| Alanine, 3-cyano- | SL Golm | 0 | 0.01 | 1 | 0.01 | 0.01 | 0.07 |
| Asparagine | ES | 0 | 0.01 | 1 | 0 | 0.01 | 0.16 |
| Aspartate | ES | 0.05 | 0.25 | 1 | 0.03 | 0.04 | 0.32 |
| GABA | ES | 0.04 | 0.18 | 1 | 0.02 | 0.04 | 0.16 |
| Glutamate | ES | 0.16 | 1 | 0.69 | 0.19 | 0.3 | 0.5 |
| Glutamine | ES | 0.01 | 0.51 | 1 | 0.02 | 0.06 | 0.17 |
| Glycine | ES | 0.35 | 0.52 | 1 | 0.48 | 0.44 | 0.6 |
| Isoleucine | ES | 0.03 | 0.12 | 1 | 0.07 | 0.1 | 0.21 |
| Leucine | ES | 0.71 | 0.69 | 1 | 0.61 | 0.88 | 0.77 |
| Lysine | ES | 0.03 | 0.31 | 1 | 0.03 | 0.07 | 0.16 |
| Pyroglutamate | ES | 0.07 | 1 | 0.86 | 0.08 | 0.27 | 0.52 |
| Proline | ES | 0.12 | 0.11 | 1 | 0.05 | 0.08 | 0.13 |
| Serine | ES | 0.01 | 0.07 | 1 | 0.02 | 0.06 | 0.05 |
| Threonine | ES | 0.03 | 0.2 | 1 | 0.09 | 0.14 | 0.14 |
| Tyrosine | ES | 0.03 | 0.18 | 1 | 0.05 | 0.06 | 0.14 |
| Valine | ES | 0.04 | 0.24 | 1 | 0.09 | 0.17 | 0.3 |
| **Organic acids** | | | | | | | |
| Citrate | ES | 0.25 | 0.29 | 0.66 | 0.36 | 0.33 | 1 |
| Fumarate | ES | 0.3 | 0.21 | 0.09 | 0.59 | 0.66 | 1 |
| Glycerate | ES | 0.49 | 0.63 | 1 | 0.38 | 0.35 | 0.47 |
| 2-Imidazolidone-4-carboxylate | SL Golm | 0.02 | 0.04 | 1 | 0.03 | 0.07 | 0.45 |
| Lactate | ES | 0.75 | 0.73 | 0.86 | 0.7 | 1 | 0.87 |
| Malate | ES | 0.34 | 0.21 | 0.07 | 0.59 | 0.44 | 1 |
| Pentonate | SL NIST | 0.52 | 0.98 | 1 | 0.4 | 0.36 | 0.52 |
| Succinate | ES | 0.15 | 0.12 | 0.03 | 0.64 | 0.71 | 1 |
| Threonate | ES | 0.34 | 1 | 0.8 | 0.38 | 0.56 | 0.91 |
| **Others** | | | | | | | |
| Ethanolamine | ES | 0.75 | 0.33 | 1 | 0.26 | 0.16 | 0.19 |
| Glycerol | ES | 0.82 | 0.7 | 1 | 0.64 | 0.55 | 0.61 |
| Hexitol | SL NIST | 0.59 | 0.55 | 1 | 0.36 | 0.21 | 0.27 |
| myo-Inositol | ES | 0.31 | 0.32 | 1 | 0.36 | 0.26 | 0.14 |
| Pentol | SL NIST | 0.69 | 1 | 1 | 0.71 | 0.57 | 0.68 |
| Phosphate | ES | 0.59 | 1 | 0.78 | 0.36 | 0.3 | 0.59 |
| **Sugars** | | | | | | | |
| Deoxyhexose | SL Golm | 0.53 | 0.61 | 1 | 0.37 | 0.29 | 0.53 |
| Fructose | ES | 0.52 | 0.5 | 1 | 0.42 | 0.28 | 0.4 |
| Glucose | ES | 0.27 | 0.26 | 1 | 0.18 | 0.15 | 0.29 |
| Hexoaldose | SL NIST | 0.18 | 0.16 | 1 | 0.11 | 0.09 | 0.17 |

| | | | | | | | |
|---|---|---|---|---|---|---|---|
| Hexoketose | SL NIST | 0.86 | 1 | 0.72 | 0.58 | 0.34 | 0.39 |
| Pentose 1 | SL NIST | 0.29 | 0.73 | 1 | 0.21 | 0.27 | 0.62 |
| Pentose 2 | SL NIST | 0.21 | 0.45 | 1 | 0.13 | 0.16 | 0.41 |
| Pentose 3 | SL NIST | 0.39 | 0.44 | 1 | 0.19 | 0.17 | 0.71 |
| Sucrose | ES | 1 | 0.19 | 0.18 | 0.62 | 0.17 | 0.15 |

Values for each compound were expressed relative to the highest values, which was set as 1. Low = 0.4 mM, Medium = 2.0 mM, High = 8.0 mM, ES = calibrated standard, SL Golm = Golm Metabolome Database, SL NIST = NIST spectral library 2.0f, n= 8 per treatment.

**Table S3:** Metabolite profile of *Populus* x *canescens* xylem sap, identified by targeted (GC-MS, LC-MS) and non-targeted approaches (LC Qtof MS).

| **Metabolite** | **Metabolite identified via** | **Observed by** |
|---|---|---|
| 12-Oxo-phytodienoic acid | MS/MS (LC-MS) | 3 |
| 1-O-(4-Coumaroyl)-β-D-glucose | MS/MS (LC-MS) | 2 |
| Dihydroxybenzoate-glucoside | MS/MS (LC-MS) | 2 |
| 2-Amino-6-hydroxypurine | MS/MS (LC-MS) | 2 |
| 2-Aminobutyric acid | GOLM DB (GC-MS) | 1 |
| 2-Hydroxyglutaric acid | MS/MS (LC-MS) | 2 |
| 2-Imidazolidone-4-carboxylate | GOLM DB (GC-MS | 1 |
| 2-*O*-acetylsalicin | MS/MS (LC-MS) | 2 |
| 3-Cyano-alanine | GOLM DB (GC-MS) | 1 |
| Dihydroxybenzoic acid | MS/MS (LC-MS) | 2 |
| 4-Aminobutyric acid | GOLM DB (GC-MS) | 1 |
| 4-Hydroxybenzoate-glucoside | MS/MS (LC-MS) | 2 |
| Methylbenzoic acid | MS/MS (LC-MS) | 2 |
| Hydroxyconiferyl alcohol derivative | MS/MS (LC-MS) | 2 |
| 7-O-Methylnaringenin-7-furanosyl-(1-6)- | MS/MS (LC-MS) | 2 |
| Abscisic acid | MS/MS (LC-MS) | 3 |
| Alanine | GOLM DB (GC-MS) | 1 |
| Arbutin (*hydroquinone glucoside*) | MS/MS (LC-MS) | 2 |
| Aromadendrin-7-*O*-glucoside | MS/MS (LC-MS) | 2 |
| Asparagine | MS/MS (LC-MS) | 1 |
| Aspartic acid | GOLM DB (GC-MS) | 1 |
| Blumenol A | MS/MS (LC-MS) | 2 |
| Caffeic acid 3-glucoside | MS/MS (LC-MS) | 2 |
| Catechin | MS/MS (LC-MS) | 2 |
| Citric acid | MS/MS (LC-MS) | 1 |
| Coniferin | MS/MS (LC-MS) | 2 |
| Coniferyl aldehyde-5-hydroxyconiferyl aldehyde | MS/MS (LC-MS) | 2 |
| Coumaric acid | MS/MS (LC-MS) | 2 |
| Coumaroyl salicin (*Trichocarposide*) | MS/MS (LC-MS) | 2 |
| Desoxyhexose | GOLM DB (GC-MS) | 1 |
| D-Glucose | MS/MS (LC-MS) | 1 |
| Ethanolamine | GOLM DB (GC-MS) | 1 |
| Fructose | MS/MS (LC-MS) | 1 |
| Fumaric acid | MS/MS (LC-MS) | 1 |
| G (8-*O*-4)G’ / G(8-*O*-4)5H | MS/MS (LC-MS) | 2 |
| G (t8-*O*-4) G | MS/MS (LC-MS) | 2 |
| G(8-5)FA (*Glycosmisic acid*) | MS/MS (LC-MS) | 2 |
| G(8-*O*-4)G' | MS/MS (LC-MS) | 2 |
| G(8–*O*–4)G(8–*O*–4)S(8–5)G' | MS/MS (LC-MS) | 2 |
| G(8–*O*–4)G(8–*O*–4)S(8–8)S | MS/MS (LC-MS) | 2 |
| G(8–*O*–4)S | MS/MS (LC-MS) | 2 |
| G(8–*O*–4)S(8–5)G' | MS/MS (LC-MS) | 2 |
| G(8-*O*-4)S(8-8)S(8-*O*-4)G | MS/MS (LC-MS) | 2 |
| G(8–*O*–4)S(8–*O*–4)S(8–8)S | MS/MS (LC-MS) | 2 |

| | | |
|---|---|---|
| G(8–*O*–4)S(8–*O*–4)X(8–*O*–4)X | MS/MS (LC-MS) | 2 |
| G(8–*O*–4)X | MS/MS (LC-MS) | 2 |
| G(8–*O*–4)X(8–*O*–4)X | MS/MS (LC-MS) | 2 |
| Galloyl-glucoside | MS/MS (LC-MS) | 2 |
| Glyceric acid | GOLM DB (GC-MS) | 1 |
| Glycerol | GOLM DB (GC-MS) | 1 |
| Glycine | GOLM DB (GC-MS) | 1 |
| Grandidentatin like | MS/MS (LC-MS) | 2 |
| Grandidentatin | MS/MS (LC-MS) | 2 |
| HCH(like)-glucoside | MS/MS (LC-MS) | 2 |
| HCH-salicortin | MS/MS (LC-MS) | 2 |
| Hesperetin 7-*O*-glucoside | MS/MS (LC-MS) | 2 |
| Hexitol | NIST DB (GC-MS) | 1 |
| Hexoaldose | GOLM DB (GC-MS) | 1 |
| Hexoketose | GOLM DB (GC-MS) | 1 |
| H-Ile-Ile-OH | MS/MS (LC-MS) | 2 |
| H-Ile-Phe-OHS | MS/MS (LC-MS) | 2 |
| H-Ile-Val-OH | MS/MS (LC-MS) | 2 |
| H-Val-Phe-OH | MS/MS (LC-MS) | 2 |
| Hydroxyquinol / 5-Trihydroxybenzene | MS/MS (LC-MS) | 1 |
| *Iso*-leucine | MS/MS (LC-MS) | 1 |
| *Iso*-Salicin | MS/MS (LC-MS) | 2 |
| Jasmonic acid | MS/MS (LC-MS) | 3 |
| Jasmonoyl-Isoleucine | MS/MS (LC-MS) | 3 |
| Jasmonoyl-Leucine | MS/MS (LC-MS) | 3 |
| Lactic acid | GOLM DB (GC-MS) | 1 |
| L-Glutamate | MS/MS (LC-MS) | 1 |
| L-Glutamine | MS/MS (LC-MS) | 1 |
| L-Isoleucine | MS/MS (LC-MS) | 1 |
| L-Leucine | MS/MS (LC-MS) | 1 |
| L-Proline | MS/MS (LC-MS) | 1 |
| L-Threonine | MS/MS (LC-MS) | 1 |
| L-Valine | MS/MS (LC-MS) | 1 |
| Lysine (4 TMS) | GOLM DB (GC-MS) | 1 |
| Malate | MS/MS (LC-MS) | 1 |
| Malic acid | MS/MS (LC-MS) | 1 |
| Matairesinol | MS/MS (LC-MS) | 2 |
| Methyl jasmonate | MS/MS (LC-MS) | 2 |
| *myo*-Inositol | GOLM DB (GC-MS) | 1 |
| *N*-hydoxypipecolic acid | MS/MS (LC-MS) | 3 |
| Nicotinate D-ribonucleoside | MS/MS (LC-MS) | 2 |
| *p*-Aminobenzoic acid | MS/MS (LC-MS) | 2 |
| Pentol | NIST DB (GC-MS) | 1 |
| Pentonate | NIST DB (GC-MS) | 1 |
| Pentose 1 | GOLM DB (GC-MS) | 1 |
| Pentose 2 | GOLM DB (GC-MS) | 1 |
| Pentose 3 | GOLM DB (GC-MS) | 1 |

| | | |
|---|---|---|
| Phosphoric acid | GOLM DB (GC-MS) | 1 |
| Pipecolic acid | MS/MS (LC-MS) | 2, 3 |
| Plastoquinol-1 | MS/MS (LC-MS) | 2 |
| Populin | MS/MS (LC-MS) | 2 |
| Porphobilinogen | MS/MS (LC-MS) | 2 |
| Protochlorophyllide | MS/MS (LC-MS) | 2 |
| Quinic acid | MS/MS (LC-MS) | 1 |
| S(8-*O*-4)Esculetin | MS/MS (LC-MS) | 2 |
| S(8-*O*-4)S(8-5)G | MS/MS (LC-MS) | 2 |
| Salicin | MS/MS (LC-MS) | 2 |
| Salicortin | MS/MS (LC-MS) | 2 |
| Salicyl alcohol | MS/MS (LC-MS) | 2 |
| Salicylic acid | MS/MS (LC-MS) | 2, 3 |
| Salicylic acid-glucoside | MS/MS (LC-MS) | 2 |
| Salicyloylsalicin | MS/MS (LC-MS) | 2 |
| Salicyloylsalicin-C7H12O7 | MS/MS (LC-MS) | 2 |
| Salirepin | MS/MS (LC-MS) | 2 |
| Scopoletin-C5H9O5 | MS/MS (LC-MS) | 2 |
| Scopoletin-glucoside | MS/MS (LC-MS) | 2 |
| Serine | GOLM DB (GC-MS) | 1 |
| Succinic acid | MS/MS (LC-MS) | 1 |
| Sucrose | MS/MS (LC-MS) | 1 |
| Syringic acid-glucoside | MS/MS (LC-MS) | 2 |
| Threonic acid | MS/MS (LC-MS) | 1 |
| Tremulacin | MS/MS (LC-MS) | 2 |
| Tyrosine | MS/MS (LC-MS) | 1 |
| Vanillic acid-glucoside | MS/MS (LC-MS) | 2 |
| Xylulose | MS/MS (LC-MS) | 1 |
| β-Alanine | GOLM DB (GC-MS) | 1 |
| γ-Glutamyl-β-cyanoalanine | MS/MS (LC-MS) | 2 |
| γ-Glutamyl-γ-aminobutyraldehyde | MS/MS (LC-MS) | 2 |

1, Targeted primary metabolite analysis by GC-MS; 2, Non-targeted metabolite fingerprinting by LC Qtof MS, followed by HRMS/MS fragmentation analyses; 3, Targeted phytohormone analysis by LC-MS

**Table S4**: Composition of the nutrient solutions used to irrigate *Populus* x *canescens* during the experiment. The concentrations are in mM.

| **Compound** | **$NH_4$** | | | **$NO_3$** | | |
|---|---|---|---|---|---|---|
| | **L** | **M** | **H** | **L** | **M** | **H** |
| $NH_4Cl$ | 0.4 | 2 | 8 | 0 | 0 | 0 |
| $KNO_3$ | 0 | 0 | 0 | 0.4 | 2 | 8 |
| KCl | 4 | 4 | 4 | 3.6 | 2 | 0 |
| $CaCl_2$ x $2H_2O$ | 2 | 2 | 2 | 2 | 2 | 0 |
| $Ca(NO_3)_2$ x $4H_2O$ | 0 | 0 | 0 | 0 | 0 | 2 |
| $MgSO_4$ x $7H_2O$ | 0.300227199 | | | | | |
| $KH_2PO_4$ | 0.599897127 | | | | | |
| $K_2HPO_4$ | 0.041337498 | | | | | |
| $H_3BO_3$ | 0.009995148 | | | | | |
| $MnSO_4$ x $H_2O$ | 0.001999763 | | | | | |
| $Na_2MoO_4$ x $2H_2O$ | 0.00699318 | | | | | |
| $CoSO_4$ x $7H_2O$ | 3.98435E-05 | | | | | |
| $ZnSO_4$ x $7H_2O$ | 0.00020032 | | | | | |
| $CuSO_4$ x $7H_2O$ | 0.000128164 | | | | | |
| EDTA - Fe x $3H_2O$ | 0.008717644 | | | | | |

**Table S5**: Mass transitions and optimized parameters for detection of phytohormones by mass spectrometry. RT is the retention time. The ionization mode depicts the polarity of the nanoESI source. Q1 and Q3 are the parent and product ion, respectively. CP, EP, CE and CXP indicate the declustering potential, entrance potential, collision energy and cell exit potential for each phytohormone.

| **Phytohormone** | **RT [min]** | **Ionization mode** | **Q1 [*m/z*]** | **Q3 [*m/z*]** | **DP [V]** | **EP [V]** | **CE [V]** | **CXP [V]** |
|---|---|---|---|---|---|---|---|---|
| SA | 2.0 | Negative | 137 | 93 | -25 | -6 | -20 | -10 |
| SAG | 1.0 | Negative | 299 | 137 | -30 | -4 | -18 | -2 |
| JA-Ile | 5.0 | Negative | 322 | 130 | -45 | -5 | -28 | -2 |
| Pip | 0.7 | Positive | 130 | 84 | 90 | 8 | 22 | 4 |
| NHP | 0.7 | Negative | 144 | 82 | -60 | -8 | -15 | -13 |
| ABA | 3.5 | Negative | 263 | 153 | -35 | -4 | -14 | -2 |
| $D_4$-SA | 2.0 | Negative | 141 | 97 | -25 | -6 | -22 | -6 |
| $^{13}C_6$-SAG | 1.0 | Negative | 305 | 137 | -30 | -4 | -18 | -2 |
| $D_3$-JA-Leu | 5.0 | Negative | 325 | 133 | -65 | -4 | -30 | -2 |
| $D_9$-Pip | 0.7 | Positive | 139 | 93 | 90 | 8 | 22 | 4 |
| $D_9$-NHP | 0.7 | Negative | 153 | 90 | -60 | -8 | -15 | -13 |

**Table S6**: Quality and processing details of RNA sequencing and library generation. RNA sequencing was carried out using a NextSeq500 (Illumina) instrument and library generation with NEBNext Ultra RNA Library Prep Kit for Illumina). *Populus* x *canescens* leaves were analyzed after three weeks feeding poplar with Low (L:0.4 mM), Medium (M: 2.0 mM) or High (H: 8.0 mM) ammonium or nitrate in the nutrient solution. RIN = RNA integrity number

| Treatment | PlantNo | RIN | Raw reads | Processed (fastP) | Mapped | Mapped (%) |
|---|---|---|---|---|---|---|
| L $NH_4^+$ | 1 | 5.7 | 13783616 | 13734565 | 12763277 | 92.93 |
| L $NH_4^+$ | 4 | 5.9 | 18551988 | 18484328 | 17256570 | 93.36 |
| L $NH_4^+$ | 5 | 5.8 | 14487052 | 14434266 | 13545914 | 93.85 |
| L $NH_4^+$ | 9 | 5.6 | 14735119 | 14677327 | 13749307 | 93.68 |
| M $NH_4^+$ | 2 | 5.5 | 12954047 | 12891433 | 9986520 | 77.47 |
| M $NH_4^+$ | 4 | 5.9 | 16377042 | 16316270 | 15260381 | 93.53 |
| M $NH_4^+$ | 7 | 5.7 | 17865955 | 17791507 | 16725784 | 94.01 |
| M $NH_4^+$ | 10 | 5.7 | 14358012 | 14305123 | 13472200 | 94.18 |
| H $NH_4^+$ | 2 | 9.2 | 18486815 | 18382349 | 17130455 | 93.19 |
| H $NH_4^+$ | 4 | 9.1 | 18465611 | 18366167 | 16819299 | 91.58 |
| H $NH_4^+$ | 6 | 8.9 | 17824503 | 17723713 | 16341968 | 92.20 |
| H $NH_4^+$ | 7 | 8.9 | 16415811 | 16327898 | 15166057 | 92.88 |
| H $NH_4^+$ | 9 | 9.2 | 19294962 | 19165705 | 17989286 | 93.86 |
| L $NO_3^-$ | 2 | 5.1 | 14353980 | 14296524 | 12860357 | 89.95 |
| L $NO_3^-$ | 7 | 5.4 | 15737311 | 15680877 | 14672981 | 93.57 |
| L $NO_3^-$ | 8 | 5.4 | 21088585 | 21009692 | 18712433 | 89.07 |
| L $NO_3^-$ | 10 | 5.2 | 19085946 | 19010101 | 17561946 | 92.38 |
| M $NO_3^-$ | 2 | 5.8 | 17404205 | 17341032 | 16174619 | 93.27 |
| M $NO_3^-$ | 4 | 5.7 | 16581387 | 16515881 | 14817873 | 89.72 |
| M $NO_3^-$ | 6 | 5.5 | 16875880 | 16811495 | 14940152 | 88.87 |
| M $NO_3^-$ | 8 | 5.7 | 14858631 | 14807888 | 13873352 | 93.69 |
| H $NO_3^-$ | 2 | 8.9 | 16038666 | 15951508 | 14735951 | 92.38 |
| H $NO_3^-$ | 3 | 9.2 | 18069065 | 17966498 | 16676325 | 92.82 |
| H $NO_3^-$ | 6 | 8.8 | 17821171 | 17717955 | 16321945 | 92.12 |
| H $NO_3^-$ | 7 | 9.1 | 18702122 | 18595466 | 17301826 | 93.04 |
| H $NO_3^-$ | 9 | 8.9 | 15211278 | 15124643 | 13957851 | 92.29 |
